# Selective localization of Mfn2 near PINK1 enable its preferential ubiquitination by Parkin on mitochondria

**DOI:** 10.1101/2021.08.25.457684

**Authors:** Marta Vranas, Yang Lu, Shafqat Rasool, Nathalie Croteau, Jonathan D. Krett, Véronique Sauvé, Kalle Gehring, Edward A. Fon, Thomas M. Durcan, Jean-François Trempe

## Abstract

Mutations in Parkin and PINK1 cause an early-onset familial Parkinson’s disease. Parkin is a RING-In-Between-RING (RBR) E3 ligase that transfers ubiquitin from an E2 enzyme to a substrate in two steps: 1) thioester intermediate formation on Parkin, and 2) acyl transfer to a substrate lysine. The process is triggered by PINK1, which phosphorylates ubiquitin on damaged mitochondria, which in turn recruits and activates Parkin. This leads to the ubiquitination of outer mitochondrial membrane proteins and clearance of the organelle. While the targets of Parkin on mitochondria are known, the factors determining substrate selectivity remain unclear. To investigate this, we examined how Parkin catalyzes ubiquitin transfer to substrates. We found that His433 in the RING2 domain catalyzes acyl transfer. In cells, mutation of His433 impairs mitophagy. In vitro ubiquitination assays with isolated mitochondria show that Mfn2 is a kinetically preferred substrate. Using proximity-ligation assays, we show that Mfn2 specifically co-localizes with PINK1 and phospho-ubiquitin in U2OS cells upon mitochondrial depolarization. We propose a model whereby ubiquitination of Mfn2 is efficient by virtue of its localization near PINK1, which leads to the recruitment and activation of Parkin via phospho-ubiquitin at these sites.

## Introduction

Mutations in the *PRKN* gene cause autosomal recessive Parkinson’s disease (PD) [1]. The Parkin protein is a RING1-In-Between-RING2 (RBR) E3 ubiquitin ligase implicated in many cellular processes, including mitochondrial quality control, innate immunity, and cellular survival pathways [2]. Activation of Parkin on mitochondria requires PINK1, a kinase whose mutations also cause autosomal recessive PD [3]. PINK1 builds up on damaged mitochondria and phosphorylates ubiquitin on Ser65; phospho-ubiquitin (pUb) in turn recruits and activates Parkin on the damaged organelle [4-10]. Parkin then ubiquitinates proteins on the outer mitochondrial membrane (OMM) [11-13]. This induces either the recruitment of autophagy receptors for the initiation of mitophagy, or the formation of mitochondria-derived vesicles (MDVs) [14-16].

RBR E3 ligases function through a two-step mechanism (Figure 1(a)), whereby an acceptor cysteine in the RING2 catalytic domain forms a thioester intermediate with ubiquitin, followed by its transfer to an amino group to form an isopeptide bond [17]. As most PD mutations in Parkin either abrogate protein translation or inactivate its biochemical activity (e.g. the C431F mutation), its biochemical activity is neuroprotective. It is thus crucial to understand how this E3 ligase is activated by PINK1 and to elucidate the mechanism by which it transfers ubiquitin to a substrate.

**Figure 1.**
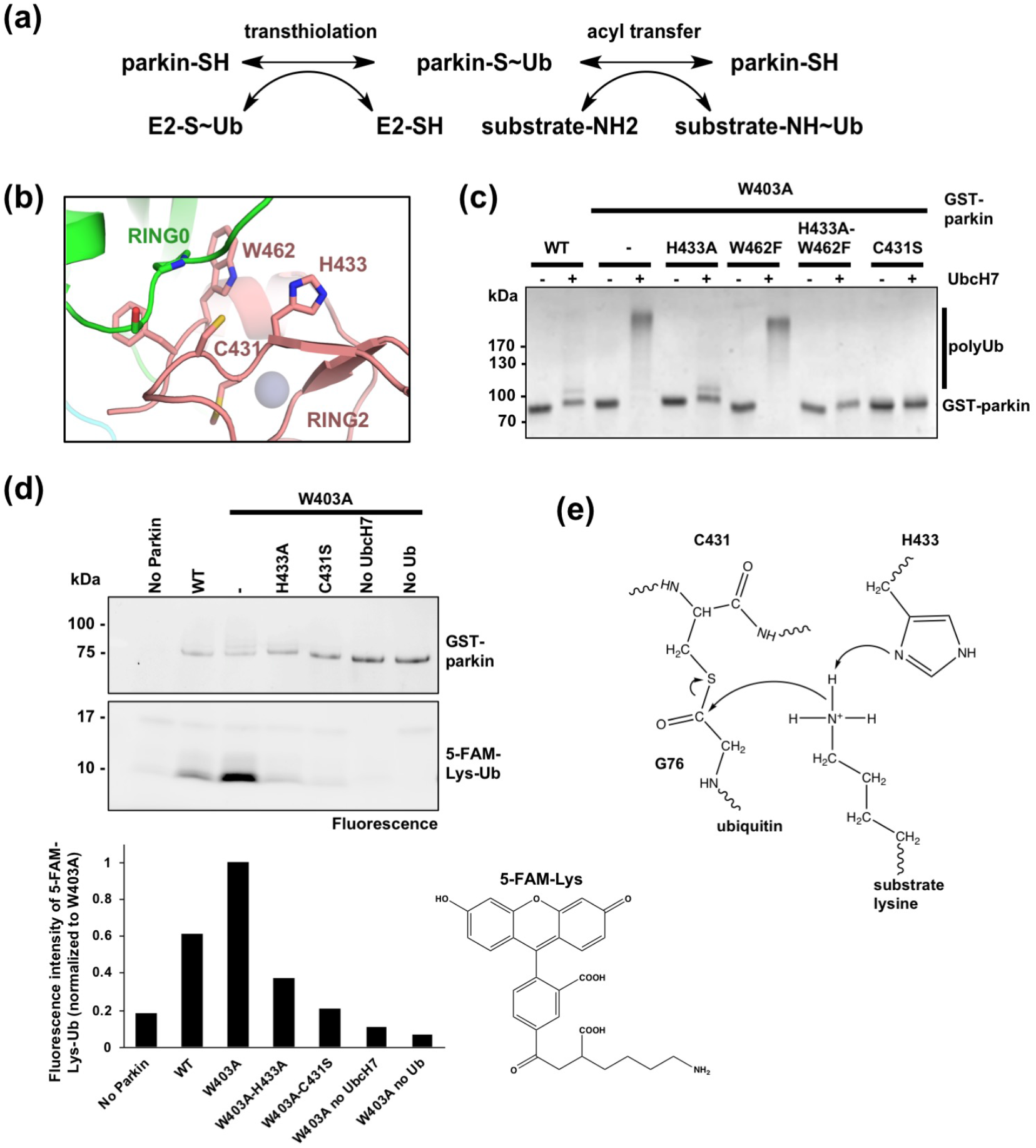
Parkin His433 contributes to the catalysis of Ub transfer. **(a)** Two-step reaction mechanism for substrate ubiquitination by parkin. **(b)** Parkin RING0 and RING2 interface with highlighted active site residues Cys431, His433 and Trp462. **(c)** Coomassie Brilliant Blue (CBB)-stained SDS-PAGE of autoubiquitination assay products for GST-Parkin with different active site mutations. **(d)** Ubiquitination reactions with GST-parkin mutants and fluorescein Lys (5-FAM-Lys). Products were resolved by SDS-PAGE and visualized by fluorescence. Quantification of the 5-FAM-Lys-Ubiquitin adduct is shown normalized to W403A. **(e)** Model for the catalysis of isopeptide bond formation by parkin: His433 retrieves a proton from the side-chain amino group of a lysine, which performs a nucleophilic attack on the Cys431∼Ub thioester bond.

The crystal structure of full-length Parkin in its *apo*, non-phosphorylated form, revealed a network of auto-inhibitory interactions [18-20]. The RING0 domain occludes the ubiquitin acceptor site Cys431 in RING2 and the Repressor Element of Parkin (REP) binds RING1 and blocks its E2-binding site. The ubiquitin-like (Ubl) domain also binds to a site in RING1 adjacent to the E2-binding site [18], in agreement with a previous report showing that the Ubl also inhibits Parkin’s ligase activity [21]. Following PINK1 activation, pUb binds Parkin via its RING1-IBR domains, which induces the dissociation of the Ubl domain [22-25] and its subsequent phosphorylation by PINK1 on Ser65 [26, 27]. Recently, the structure of insect phospho-Parkin in complex with pUb and UbcH7 [28] and the structure of human phospho-Parkin bound to pUb [29] were solved by X-ray crystallography. Both structures revealed that the phosphorylated Ubl domain binds to the RING0 domain, which induces the dissociation of the REP and RING2 domains and E2 binding on RING1 [28, 29]. The catalytic RING2 is thus free and can bind to the E2∼Ub conjugate complex to allow the transfer of ubiquitin onto Cys431 linked by a thioester bond. Kinetically, obstruction of the E2-binding site by the REP and occlusion of the active site Cys431 by RING0, as well as the inability to observe the formation on an oxyester-linked Ub on the parkin RBR C431S mutant [17], suggest that transthiolation is the rate-limiting step in parkin.

While structural and biochemical work provided much insight into the conformational changes that enable thioester transfer from the E2 to Parkin, the subsequent transfer of ubiquitin to a substrate remains far less understood. Formation of an isopeptide bond by HECT or RBR ligases involves the transfer of the ubiquitin carboxy terminus from a cysteine thiol on the ligase to a substrate amino group, typically the side-chain of a lysine [30, 31]. However, the pKa of these amino groups is typically between 9.5 and 11.5 and are thus mostly protonated at near-neutral physiological pH [32]. For the nucleophilic attack reaction to proceed, the primary amino group needs to be deprotonated. The crystal structures of *apo* Parkin [18-20] and the solution NMR structure of the RING2 domain from fly Parkin [33] revealed that His433, a residue conserved in other RBR ligases, could act as a general base to catalyze acyl transfer. His433 is adjacent to Cys431, and NMR analysis showed that this histidine is deprotonated at neutral pH and could serve as a proton acceptor [33]. The mutation H433A indeed reduced autoubiquitination activity [18-20, 33]. Mutation of the corresponding histidine in the RBR protein HOIP (His887) traps the HOIP∼Ub thioester conjugate, which is consistent with a general base role for this histidine [34]. However, this has not been demonstrated for Parkin, and the pH dependence of Parkin H433A and H433N mutants’ reactivity with ubiquitin vinyl sulfone (UbVS) suggests that it also modulates the pKa of the thiol group on Cys431, which could affect thioester transfer [20].

By contrast with HOIP, which selectively synthesizes linear polyubiquitin chains, Parkin is less selective and can conjugate all lysine side-chains on ubiquitin, primarily Lys6, Lys11, Lys48 and Lys63 [35, 36]. It can also conjugate ubiquitin to lysine side-chains on itself as well as vast array of intracellular proteins, mostly proteins located at the OMM following mitochondrial depolarization [11, 12, 35, 36]. While this would seem to indicate that Parkin does not display any specificity, recent studies suggest otherwise. Quantitative proteomics and dynamic analysis in HeLa cells and neurons with low (endogenous) levels of Parkin show that different sites on OMM proteins build up at different rates in the first hour following depolarization with oligomycin / antimycin A (OA) [37]. The most abundant ubiquitinated sites are found in VDAC1-3, the most abundant OMM proteins, followed by Mfn2, CISD1, RHOT1, FAF2, HK1, TOMM20 and TOMM70. In another study from our own group, we found that Mfn1 and Mfn2 were ubiquitinated and degraded more rapidly than the other substrates following addition of the protonophore carbonyl cyanide m-chlorophenyl hydrazone (CCCP) [38]. Others had also previously observed that Mfn1/Mfn2 are extensively ubiquitinated by Parkin [12, 13, 39].

Here, we exploit recent structural and proteomics findings to understand the biochemical basis for substrate ubiquitination by Parkin. We find that His433 acts as a base that accelerates the acyl transfer step, but is minimally involved in thioester transfer from the E2∼Ub complex. Mutation of His433 had a mild effect on mitochondrial Parkin recruitment, but a stronger effect in mitophagy. Ubiquitination assays with isolated mitochondria showed that at low Parkin concentration, Mfn2 is efficiently ubiquitinated, whereas other OMM proteins are not. In this assay, mutation of His433 only mildly affects Mfn2 ubiquitination, which is consistent with a model where Mfn2 is optimally positioned to receive ubiquitin from Parkin following activation, and thus for which the acyl transfer step is not the rate-limiting step. Using proximity-ligation assays, we demonstrate that PINK1 localizes in proximity to Mfn2, but not with other substrates, which explains why Parkin rapidly ubiquitinates Mfn2.

## Results

### Parkin His433 contributes to the catalysis of Ub transfer to a Lys ε amino group

The structure of the RING2 domain shows that the active site Cys431 is surrounded by two sidechains with acid/base chemical groups that could affect the pKa of an incoming substrate amino group: His433 and Trp462 (Figure 1(b)). To test the role played by these two residues on acyl transfer catalysis, we mutated them to Ala and Phe, respectively, and performed *in vitro* autoubiquitination assays where we monitored the formation of polyubiquitin chains on parkin after incubation with E1, UbcH7, Ub and ATP-Mg, in the context of the hyperactive W403A mutant [18]. The results confirm that the W403A mutant autoubiquitinates to a greater extent than wild-type (WT) parkin (Figure 1(c)). When paired with a mutation on the active site that inactivates ligase activity, C431S, the double mutant W403A-C431S displays almost no activity. Crucially, the double mutant W403A-H433A has impaired ubiquitination, whereas the W462F mutation has no effect on activity. The triple mutant W403A-H433A-W462F shows less activity than the W403A-H433A mutant. This suggests that His433 plays a major role in ubiquitination, whereas Trp462 plays a minor role.

To investigate the acyl transfer of ubiquitin from Parkin to a substrate Lys ε amino group, a fluorescence lysine compound (5-FAM-Lys, Figure 1(d)) was added to the ubiquitination reaction mix. After 2h incubation, SDS-PAGE-resolved products were quantified by fluorescence at 488 nm (Figure 1(d)). A control N-terminal 5-FAM-labeled ubiquitin and reactions lacking either UbcH7 or Ub were included. The results show that 5-FAM-Lys-Ub formation is observed in WT Parkin and is enhanced by the W403A mutation. In the W403A background, the two active site mutants C431S and H433A show reduced 5-FAM-Lys-Ub formation. Based on the 3D structure of Parkin [18] and these *in vitro* biochemical assays, we propose a model for the catalysis of isopeptide bond formation by parkin whereby His433 retrieves a proton from the side-chain amino group of a lysine, which performs a nucleophilic attack on the Cys431∼Ub thioester bond (Figure 1(e)).

### Parkin His433 is not essential for transthiolation

The reduced activity of the H433A mutant could also result from an impairment in thioester transfer of Ub from the E2 to Parkin Cys431. In order to assess transthiolation directly, single-turnover assays were performed. We used ubiquitin-loaded E2 (UbcH7∼Ub) and monitored disappearance of UbcH7∼Ub and autoubiquitination of Parkin through time. The products were resolved by SDS-PAGE and confirm that the hyperactive W403A mutant discharges UbcH7∼Ub faster than WT (compare 2 and 5 min time points), whereas mutation of Cys431 abolishes discharging (Figure 2(a) and Figure S1(a)). The W403A-H433A mutant discharges at the same rate as W403A, but fails to autoubiquitinate, which is consistent with His433 being required for acyl transfer and not transthiolation. The structure of auto-inhibited Parkin also shows that the side-chain of Glu444 forms a polar interaction with His433 and thus could affect acyl transfer catalysis. We thus tested the mutation E444Q, a naturally occurring variant that does not impair mitophagy [40]. The double mutant W403A-E444Q discharges UbcH7∼Ub similarly to W403A, but can still form polyubiquitin chains, suggesting it does not play a major role in either reaction.

**Figure 2.**
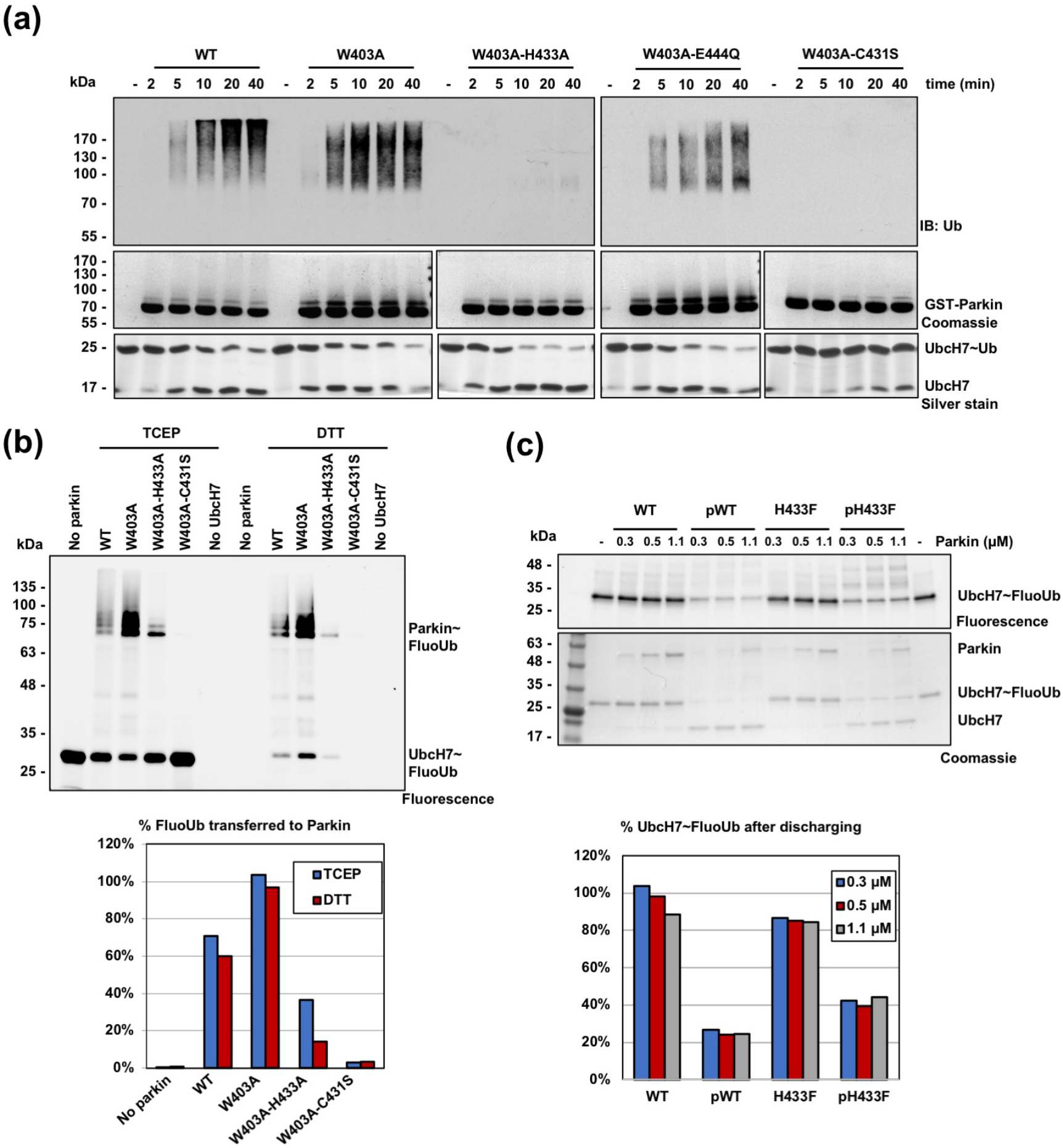
Parkin His433 is not essential for transthiolation. **(a)** Immunoblots (top) and CBB-stained SDS-PAGE (bottom) of UbcH7∼Ub discharge assays with GST-parkin wild-type and mutants. Reactions were stopped with sample buffer containing tris(2-carboxyethyl)phosphine (TCEP) to reduce disulfide bonds but keep thioester bonds intact. **(b)** SDS-PAGE fluorescent scan showing Ub-linkage after mixing fluorescein-labelled Ub (FluoUb), E1, ATP and UbcH7 with GST-parkin and stopping the reaction with TCEP or DTT (top). Quantification of FluoUb transferred to Parkin as a percentage of UbcH7∼FluoUb in the absence of Parkin (bottom). **(c)** Fluorescent scan and Coomassie-stained SDS-PAGE showing UbcH7∼FluoUb discharging by increasing concentrations of Parkin or phosphorylated Parkin (top). Quantification of Ub loaded UbcH7 (UbcH7∼FluoUb) after reaction (bottom).

One prediction from our model is that the H433A mutant would still be able to form a thioester Parkin∼Ub complex. To detect thioester formation in Parkin, we used fluorescently labelled ubiquitin (fluorescein-Ub, FluoUb) to monitor ubiquitinated species on Parkin. Reactions were stopped in the presence of either tris(2-carboxyethyl)phosphine (TCEP), which breaks only disulfide bonds, or dithiothreitol (DTT), which reduces both disulfide and thioester bonds. Ub linkages were observed by SDS-PAGE analysis and fluorescence imaging. The results show that UbcH7∼Ub is completely discharged by the DTT treatment, as expected. The gel also shows a mono-ubiquitinated form of Parkin W403A-H433A that is visible with TCEP and reduced by 60% upon addition of DTT, consistent with a mixture of isopeptide and thioester formation (Figure 2(b) and Figure S1(b)). By contrast, small differences are observed in Parkin-Ub levels between TCEP and DTT for WT and W403A, showing that these bands are not thioester-linked.

We next decided to investigate discharging of UbcH7∼Ub in the context of Parkin activation by phosphorylation. Parkin was phosphorylated in vitro by PINK1 as described before [26]. Discharging reaction mixes were prepared with different concentrations of inactive or activated Parkin. Here, His433 was mutated to a Phe, which has a bulky side-chain similar to His, but lacking acid/base chemical groups. Inactive, dephosphorylated Parkin does not significantly discharge UbcH7∼Ub, in both WT and H433F backgrounds. Once phosphorylated, both WT and H433F Parkin show increased UbcH7∼Ub discharging (Figure 2(c)). Altogether, the data is consistent with His433 being involved in acyl transfer of ubiquitin to a substrate lysine, but not in thioester transfer.

### Mutation of Parkin His433 affects mitophagy

Upon mitochondrial damage, PINK1-dependent Parkin activation and translocation from the cytosol to the mitochondria leads to the build-up of Ub chains and triggers mitophagy [7-10]. We thus sought to investigate the impact of ubiquitin acyl transfer impairment on Parkin recruitment and mitophagy. We used U2OS cells stably expressing GFP-parkin WT, H433F, or C431S. Cells were treated with CCCP to induce mitochondria membrane depolarization and imaged to track GFP-Parkin recruitment to Tom20-labelled mitochondria. As previously observed, the C431S mutant was completely impaired in recruitment (Figure 3(a)). By contrast, the H433F mutant was recruited to mitochondria, albeit more slowly than WT (Figure 3(a)). We indeed observe that the H433F mutant is capable of ubiquitinating OMM substrates such Mfn1/2 and VDAC in cells (Figure 3(b)), in line with the proposed positive feedback mechanism boosting parkin recruitment, but in apparent contradiction with our in vitro results. However, when we quantified cells negative for TOM20-mitochondria after 24h of CCCP treatment (mitophagy), cells expressing the mutants H433F and C431A were both significantly impaired in comparison with WT (Figure 3(c)). Furthermore, the distribution of diffuse and puncta GFP-Parkin in cells after 24h of CCCP treatment was also altered by the H433F mutation (Figure 3(d)). These data directly associate challenges in ubiquitin acyl transfer with impaired mitophagy.

**Figure 3.**
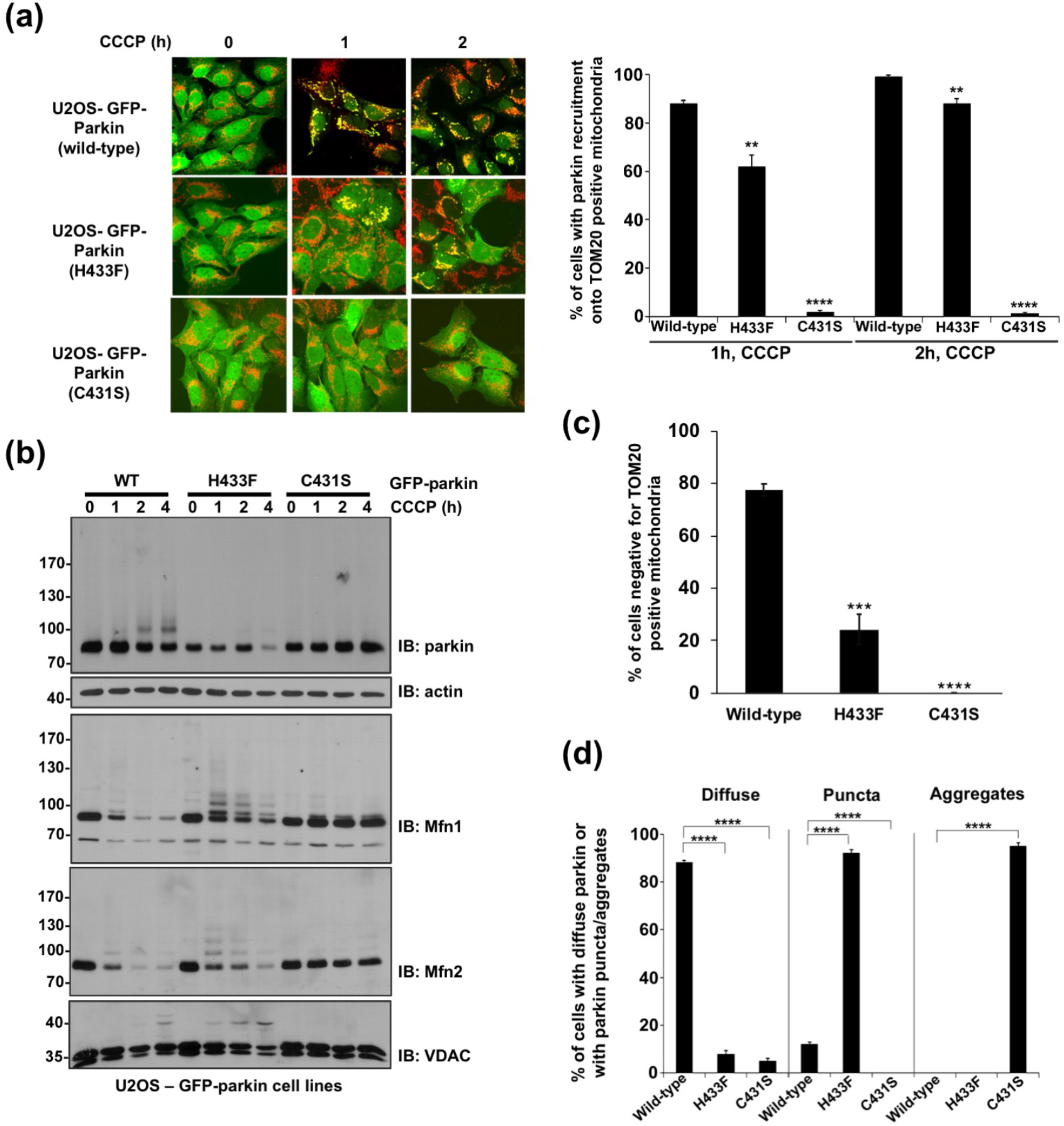
His433 is required for efficient Parkin mitochondrial recruitment. **(a)** Confocal microscopy images showing recruitment of stably expressed GFP-parkin WT, H433F or C431S to mitochondria (Tom20 antibody in red) in U2OS cells (left) and quantification of the percentage of GFP-parkin on mitochondria (right) (n=3, 2-way ANOVA, with Tukey post-test). **(b)** Western blot showing ubiquitination of mitochondrial proteins in U2OS cells after CCCP treatment. **(c)** Quantification of Tom20-negative U2OS cells expressing recombinant Parkin after 24h of CCCP treatment from immunofluorescence microscopy (100 cells analyzed per condition in 3 independent blinded experiments, 2-way ANOVA, with Bonferroni post-test). **(d)** Dispersion of GFP-Parkin in cells after 24h of CCCP treatment (n=3, 2-way ANOVA, with Tukey post-test). **p<0.01; ***p<0.001; ****p<0.0001

### Ubiquitination assays on isolated mitochondria reveals preference for Mfn2

In order to understand how substrate ubiquitination might be differentially affected by mutation of His433 in cells, we set up a series of *in organello* ubiquitination assays, a cell-free assay with isolated mitochondria previously described [26]. Briefly, HeLa cells, which lack Parkin, can be treated with CCCP to allow PINK1 build up at the OMM prior to the isolation of mitochondria. Addition of a ubiquitination mix (ATP, E1, UbcH7 and Ub) and recombinant catalytically active Parkin to isolated mitochondria is sufficient to ubiquitinate OMM substrate Mfn2. (Figure 4(a))

**Figure 4.**
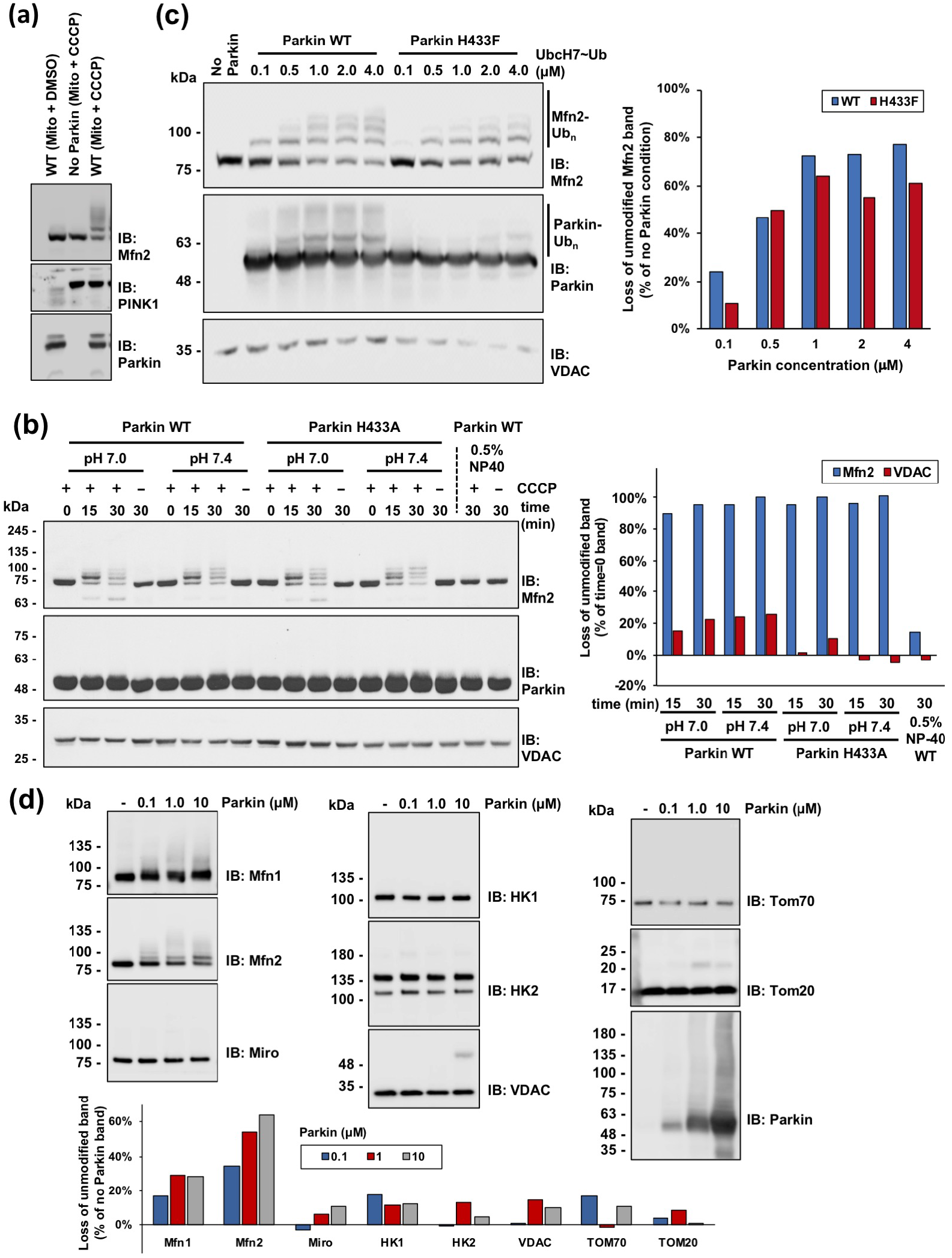
Ubiquitination assays on isolated mitochondria reveal preference for Mfn2. **(a)** Schematics of the *in organello* ubiquitination assay, where mitochondria isolated from HeLa cells, treated or not with CCCP are added to a ubiquitination mix (ATP, E1, UbcH7 and Ub) and recombinant Parkin (left). Ubiquitinated products are detected by western blot analysis (right). **(b)** Immunoblot of *in organello* ubiquitination assay testing activity of active site H433A mutation at different pH and time. Quantification of unmodified VDAC and Mfn2 bands is shown on the right. **(c)** Immunoblot of *in organello* ubiquitination assay testing activity of active site H433F mutation at different concentrations of Ub-loaded UbcH7 (UbcH7∼Ub). Quantification of unmodified band is shown on the right. **(d)** Immunoblots of *in organello* ubiquitination assays to evaluate ubiquitination levels of outer mitochondrial membrane proteins. Quantification of unmodified bands is shown below.

Reaction products were resolved by SDS-PAGE and positive ubiquitination observed by the formation of high-molecular weight bands and loss of the unmodified band. Results show that extensive ubiquitination of Mfn2 is dependent on both depolarization of mitochondria and the presence of the Parkin (Figure 4(a)). Ubiquitination of Mfn2 under these conditions is insensitive to the active site H433A mutation and proportional to the amount of Parkin in the reaction (Figure 4(b) and Figure S2(a)). While monoubiquitination of VDAC can be observed at high concentrations of Parkin, there is little loss of the unmodified band compared to Mfn2. This difference in the extent of ubiquitination between VDAC and Mfn2 suggests a kinetic advantage for Mfn2.

The amino group of lysine side chains is typically protonated at neutral pH and they are intrinsically poor nucleophiles. Given the proposed role of His433 to act as a base to deprotonate any substrates’ lysine side chain in order to facilitate the acyl transfer step, *in organello* ubiquitination assays were performed to assess the effects of deprotonation. Time-course of ubiquitination at two different pH (7.0 and 7.4) shows no difference between WT and H433A Parkin in Mfn2 ubiquitination. When we tested a wider pH range, Mfn2 ubiquitination still presented the same pattern for both WT and H433F variants (Figure S2(b)). Yet, the overall ubiquitination levels detected at high molecular weight protein bands was significantly affected when His433 was mutated. Since we have shown that mutation of His433 reduces acyl transfer but not thioester transfer, these *in organello* data imply that the rate-limiting step for Mfn2 ubiquitination is thioester transfer. However, acyl transfer is the rate-limiting step for autoubiquitination in vitro (Fig. 1 and 2), and overall ubiquitination of mitochondrial proteins (Figure S2(b)). Indeed, if we use pre-charged Ubch7∼Ub in organello, we observe a small reduction in Mfn2 ubiquitination by the H433F mutant (Figure 4(c)). Since thioester transfer is independent of the substrate, this data is pointing to Mfn2 being a kinetically preferred substrate for acyl transfer.

We next sought to evaluate ubiquitination of known OMM proteins by Parkin in *in organello* assays using recombinant WT Parkin in a concentration range from 0 to 10 μM. Products were resolved by SDS-PAGE and immunoblotted for the different proteins. Results confirm that Mfn2 is readily ubiquitinated for concentrations of Parkin as low as 0.1 μM (Figure 4(d)). Mfn1 is also ubiquitinated in a similar fashion, though its ubiquitination pattern is less prominent than Mfn2. The remaining substrates are not substantially ubiquitinated and mostly remain unmodified even at the highest Parkin concentration (10 μM) (Figure 4(d)). This confirms that Mfn2 is a kinetically preferred substrate.

One possible explanation for the observed preference for Mfn2 is that Parkin binds directly to this protein. However, addition of non-denaturating detergent NP40, which solubilizes mitochondrial membranes, abrogates Mfn2 ubiquitination even in the presence of CCCP (Figure 4(b), last 2 lanes). However, similar detergent concentrations did not affect Parkin’s E3 ubiquitin ligase activity in vitro (Figure S2(c)). Thus, membrane integrity is critical for Mfn2 ubiquitination by Parkin, which implies that Parkin does not engage with Mfn2 directly. Given that Mfn2 is conjugated to pUb in cells [38], this suggest that a critical step for substrate selectivity is labelling of substrates that are in proximity to PINK1 by pUb.

### Parkin ubiquitinates substrates conjugated to phospho-ubiquitin

To explore the model by which pUb labeling drives substrate selectivity, we set up in vitro ubiquitination assays where we used purified glutathione-S-transferase (GST) as “bait” fused to either ubiquitin (GST-Ub) or phosphorylated-Ub^Ser65^ (GST-pUb^S65^). In a series of reactions, purified Parkin or phospho-Parkin (100 nM) was incubated with a sole donor Ub source, UbcH7∼Ub (4 μM), in addition to GST-Ub or GST-pUb^S65^ (2 μM) as substrates (Figure 5(a)). In the absence of the E1 ubiquitin-activating enzyme, the GST-Ub substrates cannot be used as a donor Ub. Reactions were resolved by SDS-PAGE and products analyzed by immunoblotting. When UbcH7∼Ub was mixed with non-phosphorylated Parkin alone or in the presence of GST-Ub, little ubiquitination was observed (Figure 5(b), lanes 5, 7). When UbcH7∼Ub was mixed with phosphorylated Parkin instead, alone or in the presence of GST-Ub, autoubiquitination levels increased (Figure 5(b), lane 6,11) as expected, but no substantial GST-Ub ubiquitination was observed. When GST-pUb^S65^ is used instead, Parkin autoubiquitination is increased and critically GST-pUb^S65^ ubiquitination is detected, as observed in the appearance of a ladder in pUb and Ub blots (Figure 5(b), lane 8). Most strikingly, ubiquitination of GST-pUb^S65^ was further increased in the presence of phospho-Parkin, as observed in the Ub and pUb blots as well as loss of the unmodified GST-pUb band in the Ponceau stain (Figure 5(b), lane 12).

**Figure 5.**
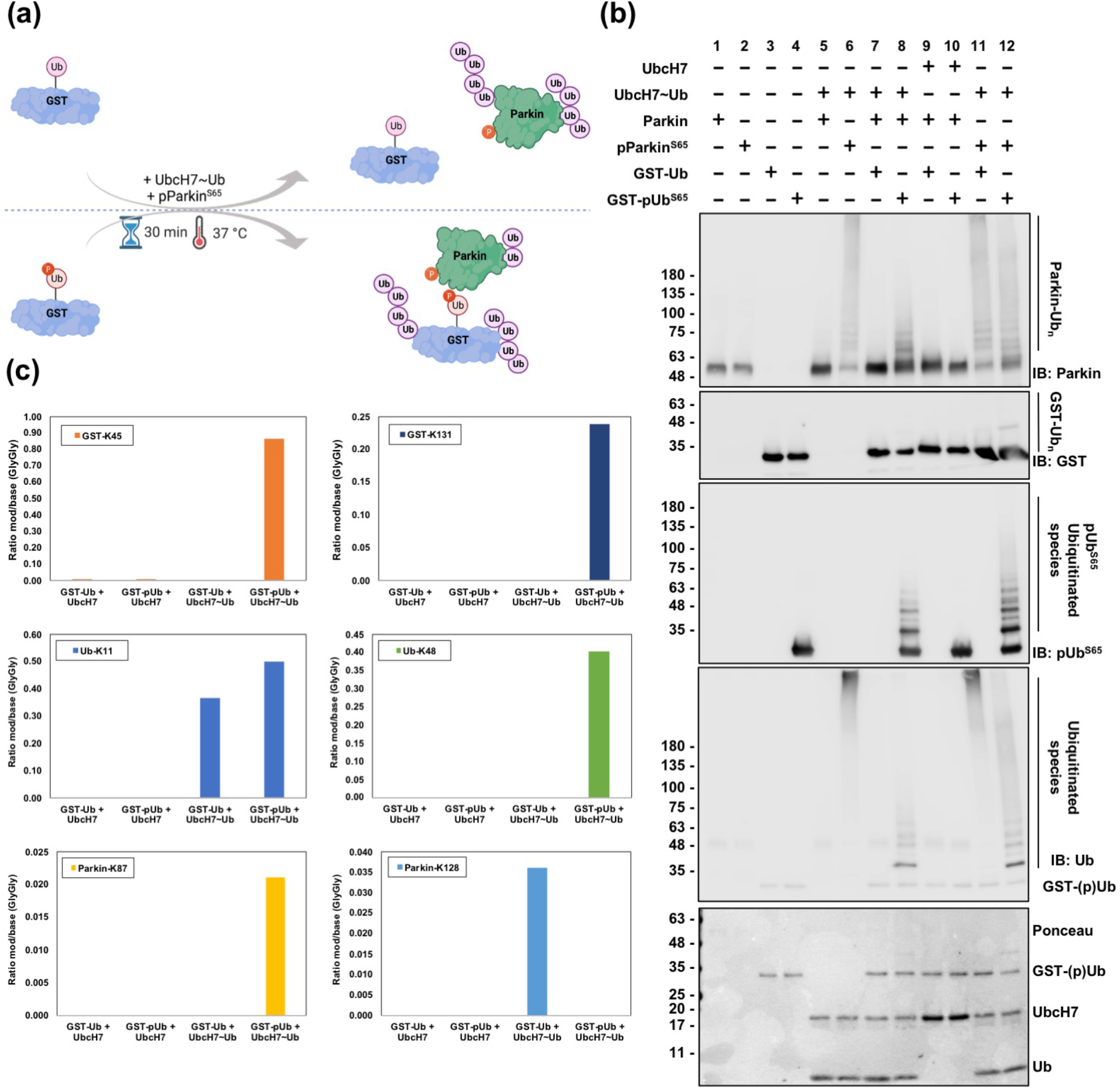
Parkin preferably ubiquitinates substrates tethered to phospho-ubiquitin. **(a)** Schematics of the ubiquitination assay where Glutathione-S-transferase (GST) fused to either ubiquitin (GST-Ub) or phosphorylated ubiquitin (GST-pUb^S65^) was used as a Ub acceptor and shown in (b). **(b)** Immunoblots of the ubiquitination assay where purified Parkin or phosphorylated Parkin (pParkin^S65^) was incubated with UbcH7∼Ub as a ubiquitin source and GST-Ub or GST-pUb^S65^ as the ubiquitin acceptor. Lanes 1 to 10 are reaction controls. Bands at around 48 kDa in lanes 1, 2, 5, 7, 9 and 10 are cross reactive bands. **(c)** Mass spectrometry analysis of the ubiquitination sites found for lanes 9 to 12 shown in (b). Selected sites in GST, Ub and Parkin molecules are shown.

To confirm and determine where GST-pUb^S65^ was ubiquitinated, and also because ubiquitination of GST likely affects the immunoreactivity of the anti-GST antibody, we analyzed the reaction products with trypsin digestion and mass spectrometry (Figure 5(c) and Figure S3(a)). Products from reactions containing UbcH7/UbcH7-Ub, Phospho-Parkin and either GST-Ub (Figure 5(b), lanes 9,11) or GST-pUb^S65^ (Figure 5(b), lanes 10, 12) were analyzed using MaxQuant to identify and quantify lysine residues modified with the GlyGly motif left after trypsin cleavage of the Ub C-terminus. Two solvent-exposed lysine residues in GST (Lys45 and Lys131) were strongly ubiquitinated in GST-pUb^S65^ and could not be detected in GST-Ub (Figure 5(c) and Figure S3(b)). Weaker sites were identified elsewhere and showed a similar pattern (Figure S3(a)). Because of its low abundance in the reaction mix, Parkin peptides were difficult to detect, but ubiquitination sites were identified on two lysine residues located in the Ubl-RING0 linker. Intriguingly, we observed differences in the distribution of Ub linkage types (Lys11, Lys48, Lys63) between GST-Ub and GST-pUb^S65^, with a massive increase in Lys48 linkage detected in the GST-pUb^S65^ sample (Figure 5(c) and Figure S3(b)). This may reflect changes in the distributivity of donor Ub, which form shorter and more widely distributed chains in the presence GST-pUb^S65^ (Figure 5(b), lane 12), and longer chains in its absence, predominantly on Parkin (Figure 5(b), lane 11). Altogether, this data supports the hypothesis that when a protein is phospho-ubiquitinated, it is predisposed to be a preferential substrate for ubiquitination by Parkin.

### PINK1 co-localizes with Mfn2 on damaged mitochondria

Given our previous observations about the role of pUb in dictating substrate selectivity and the extensive Mfn2 ubiquitination observed in cells and in organello, we hypothesize that Mfn2 localizes near PINK1, thus making Mfn2-Ub conjugates more likely to be phosphorylated. To test this idea, proximity ligation assays (PLA) were performed in U2OS cell lines [41]. PLA is an antibody-based assay that detects close contact sites between proteins by producing fluorescent signals at the sites of interaction, which can then be visualized with confocal microscopy (Figure 6(a)).

**Figure 6.**
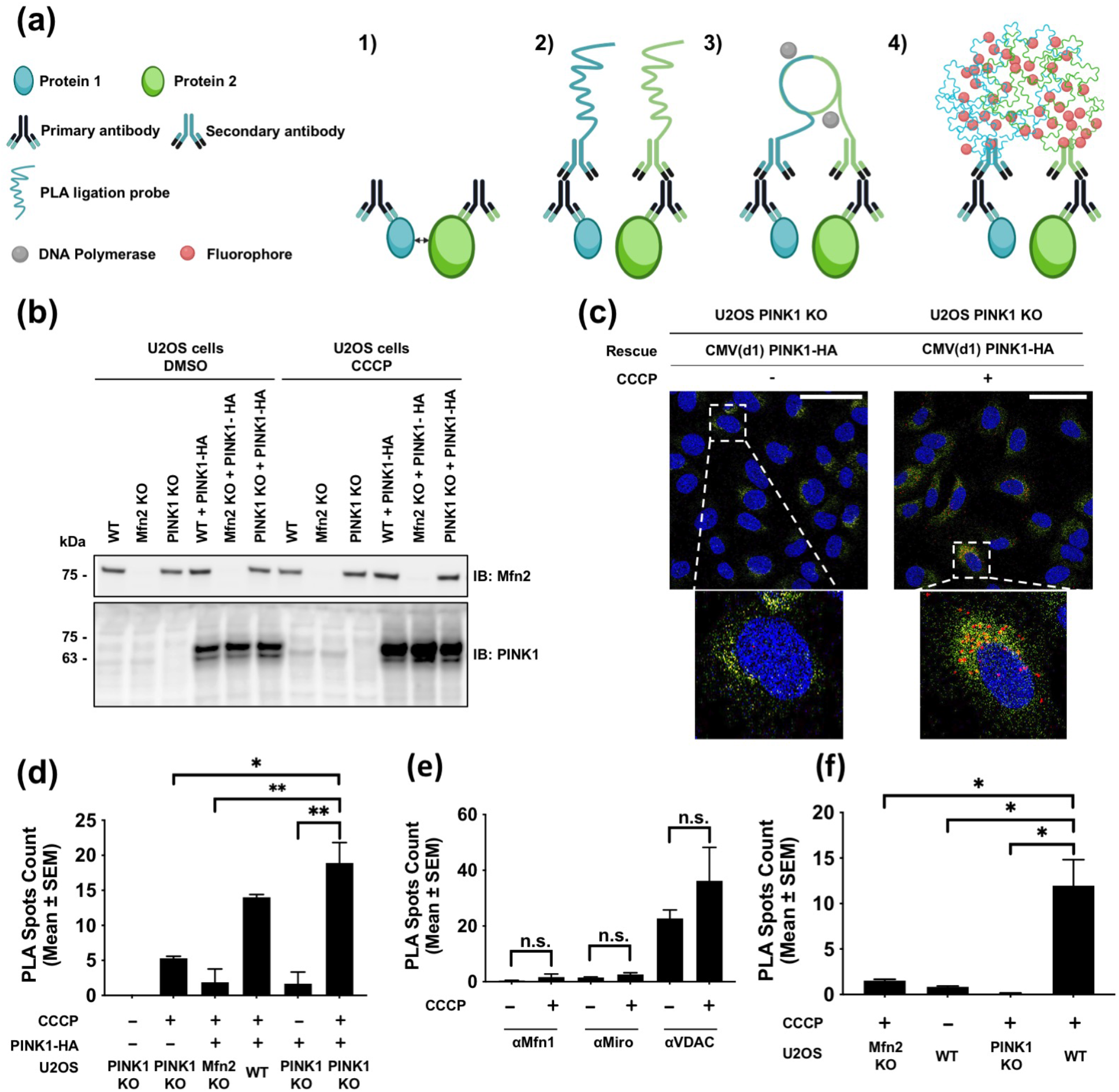
Co-localization of PINK1 and Mfn2 on damaged mitochondria using the proximity ligation assay (PLA). **(a)** Schematic overview of PLA. 1) Two primary antibodies raised in different species recognize two proteins (< 40 nm). 2) Secondary antibodies coupled with PLA ligation probes bind to the primary antibodies. 3) PLA ligation probes in proximity ligate to form a circular DNA template amplified by DNA polymerase. 4) Fluorophores hybridize to the template and amplify the fluorescent signals detectable by microscopy. **(b)** Immunoblots showing the Mfn2 and PINK1 expression levels in U2OS knockout (KO) cells. **(c)** PLA confocal microscopy images targeting Mfn2 and PINK1-HA in U2OS PINK1-KO cells. PLA spots (red) are located around the nuclei (DAPI stain, blue) and within the contour of the ER (Alexa488-coupled anti-calnexin, green). Scale bar, 10 microns. **(d-f)** Quantification of PLA spots per cell for interactions between Mfn2 and PINK1 (d), Mfn1, MIRO or VDAC, and PINK1 (e) and between Mfn2 and phospho ubiquitin (f). Data from three independent experiments was tested for significance with independent t-tests with Bonferroni corrections post-hoc test were performed. The vertical bars represent s.e.m. ** P < 0.01. *** P < 0.001 (D); n.s., not significant (e); * P < 0.05. ** P < 0.01 (f)

Because of technical challenges in labeling both endogenous PINK1 and Mfn2 with antibodies from different species, as required for PLA, a CMV(d1) promoter-driven PINK1-HA plasmid was used [42]. PINK1 expressed from this plasmid accumulates on depolarized mitochondria and generates pUb chains [27]. Experiments were performed in both WT and PINK1 KO cells, as well as Mfn2 KO cells as negative controls (Figure 6(b)). Results show that PINK1 (endogenous or ectopic) accumulates in the presence of CCCP, independently of Mfn2.

PLA experiments targeting Mfn2 and PINK1-HA (anti-HA) were first performed and the fluorescent spots corresponding to close contact sites between the two labelled proteins quantified. Results show that an average of less than 5 PLA spots per cell were counted in negative controls where either PINK1 or Mfn2 is absent (Figure 6(c) and Figure 6(d)). Few PLA spots are also observed in cells without CCCP treatment, where there is no mitochondrial depolarization causing PINK1 buildup. Conversely, a significant number of fluorescent PLA spots were observed around the nucleus and within the contour of ER are accounted for U2OS WT and PINK1 KO cells, when expressing recombinant PINK1-HA (Figure 6(c) and Figure 6(d)).

We next assessed the co-localization of PINK1-HA with other OMM Parkin substrates in U2OS PINK1 KO cells. PLA was performed with labelled Mitofusin-1 (Mfn1), voltage-dependent anion channel (VDAC), and Miro1. When targeting Mfn1 and PINK1, PLA signals are nearly undetectable in both the negative control and the CCCP-treated experimental condition (Figure 6(e) and Figure S4(b)). A similar observation is obtained when targeting Miro and PINK1 (Figure 6(e) and Figure S4(c)). A large number of PLA signals is observed when targeting VDAC and PINK1-HA, yet these are also present in the absence of CCCP (Figure 6(e) and Figure S4(d)). In sum, none of the three tested OMM proteins are located or enriched in close proximity to PINK1 upon mitochondrial depolarization.

To further consolidate our observation, we sought to assess whether Mfn2 is found in close proximity to pUb, which is the direct output of PINK1’s catalytical activity. PLA assays were performed without employing an ectopic overexpression system (i.e. transfecting exogenous PINK1-HA) to evaluate the interaction between proteins of interest at endogenous levels of PINK1 in cells. The total number of detected PLA spots show that Mfn2 is also found in proximity to pUb in cells as these signals are absent in Mfn2 KO, PINK1 KO or untreated control conditions (Figure 6(f) and Figure S4(e)). Collectively, these data strongly demonstrate that Mfn2 preferentially localizes in close proximity to PINK1 and pUb.

## Discussion

Formation of an isopeptide bond by HECT or RBR ligases involves the transfer of the ubiquitin carboxy terminus from a cysteine thiol on the ligase to a substrate amino group. As these residues are mostly protonated at physiological pH, catalysis of deprotonation will lead to efficient acyl transfer. Our data show that His433 plays an important role in the acyl transfer of Ub to substrates (Figures 1 and 2), consistent with those obtained for HOIP [34]. The structure of HOIP bound to ubiquitin (in both donor and acceptor position) shows the acceptor Ub bound to RING2, with its amino terminal Met1 group positioned adjacent to Cys885 and His887, while Gly76 from the donor Ub is bound to Cys885 (Figure 7(a)). Cys431 and His433 in the Parkin inactive structure are oriented in a similar manner, with the RING0 domain occupying the position of the acceptor Ub in HOIP. Upon phosphorylation of the Ubl, the RING2 domain dissociates from RING0, which would allow Cys431 to form a thioester bond with a donor Ub and transfer it to an acceptor substrate amino group. One key difference between HOIP and Parkin is that the RING2 domain in HOIP is followed by a zinc finger and the linear ubiquitin chain determining domain (LDD), which form contacts with the acceptor Ub and are essential for dictating chain specificity [43]. Parkin lacks such elements, which could explain its lack of substrate selectivity.

**Figure 7.**
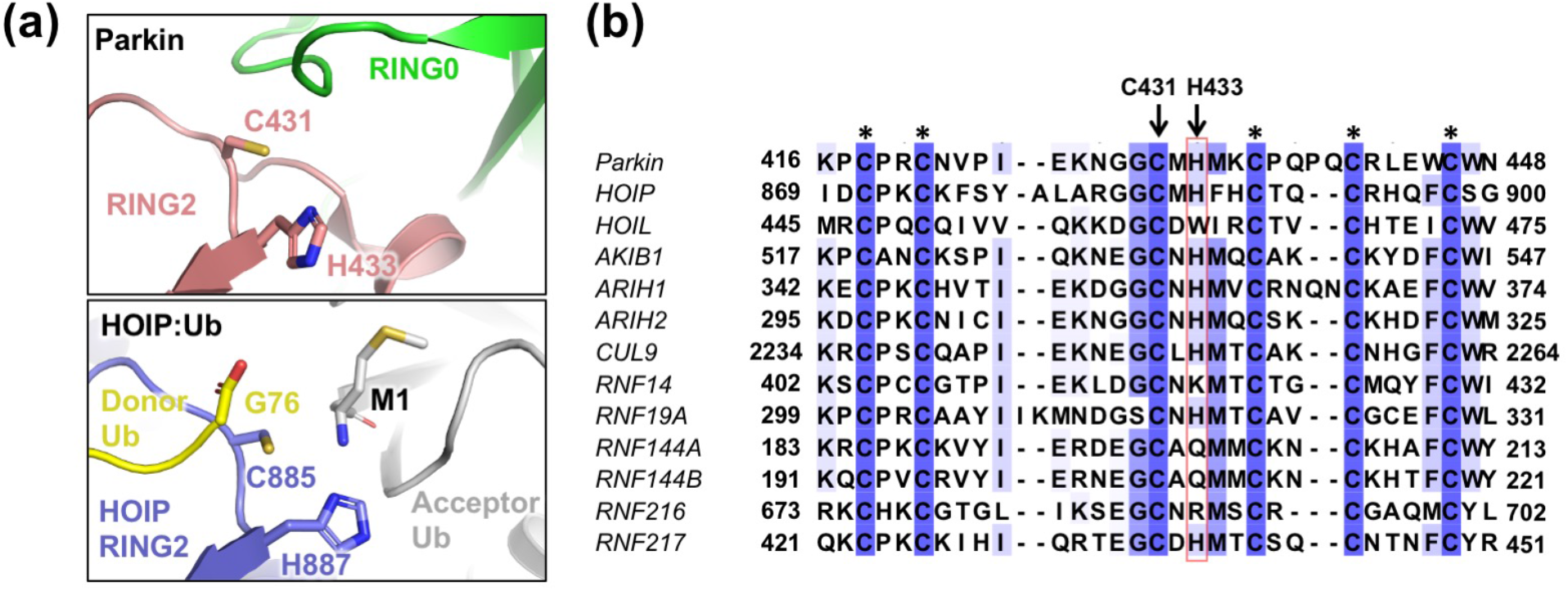
Conservation and structure of Parkin His433. **(a)** Comparison of the RING2 domains from autoinhibited parkin (pdb 4K7D) and HOIP bound to donor and acceptor ubiquitin molecules (pdb 4LJO). Met1 of the acceptor Ub is close to His887 in HOIP, which act as a general base by deprotonating the terminal amino group [33]. The C-term of the donor Ub is close to the catalytic cysteine, mimicking the Ub∼Cys885 thioester intermediate. The RING0 domain in Parkin overlaps with the acceptor Ub in the HOIP complex. **(b)** Sequence alignment of RING2 domains from different human RBR E3 ubiquitin ligases. Asterisks indicate zinc-coordinating residues.

While His887 in HOIP is important to catalyze Ub acyl transfer, a subsequent structure of HOIP in complex with a charged E2∼Ub conjugate showed that His887 also mediates hydrogen bonds with the E2 enzyme [44]. However, these interactions do not appear to be critical, since the HOIP H887A mutant can still catalyze Ub thioester transfer from an E2 [34]. While there is no structure of Parkin bound a to a charged E2∼Ub conjugate, our discharging assays (Figure 2) suggest that Parkin His433 is not critical for the interaction with the E2∼Ub conjugate.

Intriguingly, a sequence alignment of the RING2 domain from RBR ligases shows that the Parkin His433 residue is conserved, but is not found in HOIL-1, RNF14, RNF144A/B, and RNF216 (Figure 7(b)). In HOIL-1, the catalytic histidine residue is replaced by Trp462. HOIL-1 binds to HOIP and is required for activation of the latter, but HOIL-1 alone does not catalyze formation of free ubiquitin chains, contrary to the HOIP catalytic domain [45]. However, HOIL-1 is capable of generating ubiquitin chains via formation of hydroxylamine-sensitive oxyester bonds, in a reaction that is dependent on the active site Cys460 [46]. Thus, HOIL-1 has different catalytic requirements than other ligases that catalyze isopeptide bonds. RNF216 does generate polyUb chains in a Cys-dependent manner, but it’s unknown whether these chains are found on lysine residues or not [47]. However, it has an arginine residue at the position equivalent to His433, and thus perhaps RNF216 catalyzes acyl transfer in a different manner. Finally, RNF144A/B have a glutamine at the equivalent position, and can catalyze formation of Lys6, Lys11, Lys48, and Lys63 polyUb chains [48], again suggesting that there are different modes of acyl transfer catalysis.

It is important to note that other factors than His433 contribute to acyl transfer in Parkin. The local environment (electrostatics, solvent accessibility) of the RING2∼Ub conjugate may itself lower the pKa of the sidechain amino group of lysine, which will enhance its reactivity. Indeed, we observe that the H433A mutant, while impaired, can still transfer Ub to substrates in vitro, to a greater extent than the C431S mutant (Figures 1 and 2). For example, single-turnover reactions with UbcH7∼FluoUb show that the W403A-H433A mutant still forms DTT-resistant Ub adducts on Parkin (15% compared to W403A), whereas W403A-C431S does not (Figure 2(b)). Likewise, the W403A-H433A mutant is still able to form the 5-FAM-Lys-Ub adduct, with a relative activity of 20% compared to W403A after subtracting signal in the absence of Parkin (Figure 1(d)). Thus, mutation of His433 does not abolish, but rather reduces the rate of acyl transfer by 5-to 10-fold.

Furthermore, we found that the mutant H433F could still be recruited to mitochondria and induce mitophagy, albeit more slowly (Figure 3). These observations go hand in hand with the proposed positive feedback mechanism where new Parkin-made Ub-chains on mitochondria are phosphorylated by PINK1 which in turn recruit more Parkin and leads to further Ub-chain synthesis [35]. In this way, when Parkin H433F fails to efficiently transfer Ub to a substrate, it also reduces the ability to boost further ligase recruitment, impairing the ability to undergo mitophagy. On immunoblots, we noticed that substrates such as Mfn1 and Mfn2 were ubiquitinated by Parkin H433F, but compared to WT, the extent of the disappearance of the unmodified band (via degradative pathways such as the ubiquitin-proteasome pathway or mitophagy) was slightly slower (Figure 3(b)). To better understand this and separate the effect of ubiquitination from either PINK1 buildup or subsequent degradation pathways, we performed in organello ubiquitination assays on isolated mitochondria. We observed that Mfn2 ubiquitination was not affected by the His433 mutation (Figure 4(b) and Figure S2(a,b)), whereas overall ubiquitination levels were affected by the mutation (Figure S2(b)). We interpret these results by proposing that the rate-limiting step for Mfn2 ubiquitination is thioester formation (His433-independent), whereas acyl transfer is the rate-limiting step for other substrates (His433-dependent). The rate of thioester formation itself depends on a multitude of upstream steps, such as Parkin binding to pUb, phosphorylation of the Ubl and release of the REP, charging of Ub on the E2 enzyme, and binding of the charged E2∼Ub conjugate, but critically, all of those steps are independent of the substrate. This implies that acyl transfer must be faster for Mfn2 than for other substrates (more than 5/10-fold, based on estimation of residual in vitro activity by the H433A mutant), which explains why Mfn2 is the most extensively ubiquitinated substrate in our organello assay (Figure 4(d)).

Using quantitative mass spectrometry experiments, Ordureau *et al*. estimated that Mfn2 is 100 and 25 times less abundant than VDAC1 and VDAC3, respectively [37]. After one hour of oligomycin / antimycin A (OA) treatment in HeLa cells expressing Parkin, the fractional occupancy of the most abundant Lys-Gly-Gly site in VDAC3 (Lys109) is around 0.5, with around 1375 fmol of peptide per 1.5 mg of isolated mitochondria. If we extrapolate to the most kinetically favoured Mfn2 site Lys416 (55 fmol for the same amount of mitochondria, i.e. ∼25x less than VDAC3^K109^), we calculate a fractional occupancy of ∼0.5 for Mfn2^K416^, which is consistent with visual observation on immunoblots [37]. However, after only 15 min of OA treatment, the fractional occupancy has already reached ∼0.17 for Mfn2^K416^, whereas it reaches only 0.016 and 0.007 for VDAC3^K53^ and VDAC3^K109^, respectively. Those results are consistent with our own and highlight that Mfn2 is a kinetically preferred substrate of Parkin.

What, then, makes acyl transfer of Ub to Mfn2 so efficient? We reported that Mfn2 co-purifies with pUb in cells, consistent with Mfn2 being conjugated to pUb [38]. These Mfn2-pUb conjugates would recruit Parkin selectively to mitochondrial sites enriched for Mfn2 and PINK1, thus bringing Parkin in closer proximity to lysine residues in Mfn2 than in other substrates. We therefore developed an artificial GST-pUb substrate to test whether coupling to pUb conferred a kinetic advantage for Parkin ubiquitination (Figure 5). While Parkin binding to pUb does enhance its autoubiquitination activity, phosphorylation of its Ubl has a much stronger effect (Figure 5, compare lanes 6 and 8; ref [23]). Yet, phospho-Parkin was unable to ubiquitinate GST-Ub, in contrast to GST-pUb which was substantially modified by Ub chains on the GST moiety because it recruits phospho-Parkin (Figure 5, lanes 11 and 12). It is worth noting that for this assay, we used low concentrations of Parkin (200 nM), similar to those used *in organello*, in order to reduce autoubiquitination in *trans* via intermolecular interactions. Yet, we found that phospho-Parkin still autoubiquitinates, especially so in the absence of GST-pUb. This is likely due to the fact that there is a limited supply of Ub in the assay; addition of a kinetically preferred substrate such as GST-pUb drives the pool of Ub away from autoubiquitination towards GST ubiquitination.

Our model of Mfn2 ubiquitination by Parkin, based on proximity to pUb, implies that PINK1 must be in close proximity to Mfn2. Our PLA data shows that PINK1-HA and Mfn2 indeed form distinct spots in response to mitochondria depolarization (Figure 6). While we also observe slightly more spots per cell between VDAC and PINK1-HA, VDAC paralogs are 25-100 more abundant than Mfn2 (in HeLa cells and neurons [37]), and therefore we expect to see many more non-specific PLA spots, and indeed we observe as many spots in cells that were not treated with CCCP (Figure 6(e)). However, we cannot rule out non-specific antibody background for VDAC. The number of spots between Mfn2 and pUb was found to be in the same range as Mfn2 and PINK1-HA in cells treated with CCCP (compare Figure 6(d) and 6(f)), suggesting that the number of spots is limited by the low abundance of Mfn2. Yet, nearly no spots were observed in cells expressing PINK1-HA but not treated with CCCP (Figure 6(d)), even though these cells overexpress PINK1-HA at far higher levels than endogenous PINK1 (Figure 6(b)). This suggests that only a small fraction of overexpressed PINK1-HA localizes near Mfn2 in a manner similar to endogenous PINK1 in cells treated with CCCP.

The nature of the interaction between Mfn2 and PINK1 is unknown, but they are unlikely to form a strong non-covalent complex, since solubilization with detergents was sufficient to abolish Mfn2 ubiquitination by Parkin (Figure 4(b)). The two proteins more likely colocalize in the same membranous sub-compartment. Mfn2 is critical to maintain mitochondria-endoplasmic reticulum (mito-ER) contact sites [49], and Parkin-mediated ubiquitination of Mfn2 indeed regulates these contact sites physically and functionally [38, 50]. The implication of our model is that PINK1 accumulates specifically at ER-mito contact sites (Figure 8). PINK1 has already been shown to localize specifically at these contact sites during mitophagy [51], and we observed that Mfn2-PINK1 PLA spots are in proximity to the ER (Figure 6(c)). Like PINK1 itself [52], ER-mito contact sites are critical for calcium homeostasis [50] and are implicated in regulating mitochondrial fission [53]. In a recent publication, fission events linked to degradation by mitophagy were shown to take place at the periphery of mitochondria, in a process involving machineries distinct from midzone fission [54]. While Parkin was recruited specifically at the periphery, it is unknown whether PINK1 and Mfn2 also localize at the periphery. The features of PINK1 which dictate localization at distinct mitochondrial locations remain undetermined and may depend on its mRNA or specific features of its mitochondrial targeting sequence.

**Figure 8.**
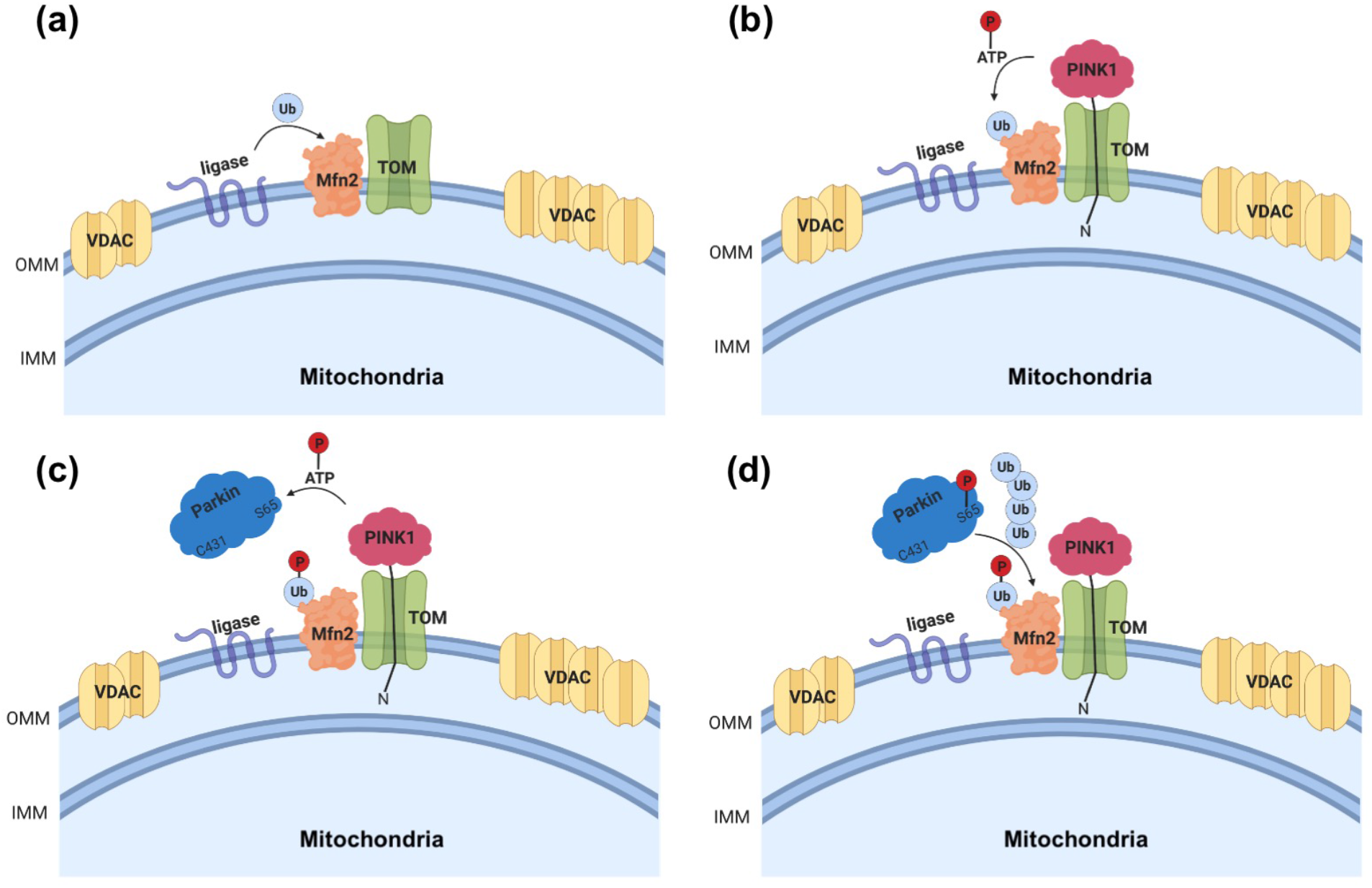
Schematic illustrating Parkin’s substrates specificity towards Mfn2 at the onset of cellular damage. **(a)** Mitochondrial E3 Ub ligases (e.g. March5) catalyze the addition of a “seed” Ub onto Mfn2. **(b)** PINK1 is recruited to the damaged OMM and uses its kinase activity to phosphorylate the existing Ub moieties on the nearby Mfn2. **(c)** The generation of pUb recruits Parkin, which becomes catalytically active upon phosphorylation at Ser65 by PINK1. **(d)** Parkin further catalyzes the addition of Ub moieties onto Mfn2, which in turn are phosphorylated by PINK1, and the feedback loop continues. Eventually, other abundant OMM substrates such as VDAC will be ubiquitinated.

We have previously shown that pUb chains build up on mitochondria in the absence of Parkin, and that binding to these so-called “pre-existing” pUb chains is required for Parkin phosphorylation and activation [26]. If PINK1 phosphorylates Ub located on OMM proteins such as Mfn2 prior to Parkin recruitment, which E3 ubiquitin ligase(s) ubiquitinate(s) Mfn2 in the first place? In addition to Parkin, three ubiquitin ligases have been shown to ubiquitinate Mfn2, namely Huwe1, Mul1, and March5. Huwe1 is a cytosolic HECT-type Ub ligase that forms Lys6-Ub chains and can ubiquitinate Mfn2 in cells [48, 55]. Mul1, also known as MAPL, is a membrane protein with a RING domain that bears ubiquitination as well as sumoylation activity, and has been reported to modulate Mfn2 [56, 57]. March5, also known as MITOL, is an integral membrane protein on the OMM that harbors a RING domain and ubiquitinates Mfn2 [58, 59]. The group of Noriyuki Matsuda knocked down all three Ub ligases and found that only March5 knockdown affected Parkin-dependent substrate ubiquitination [60]. Furthermore, they showed that March5 KO attenuated Parkin recruitment to depolarized mitochondria, whereas March5 overexpression accelerated recruitment. The group of Wade Harper later showed that KO of Mul1 or March5 did not affect pUb chain formation or substrate ubiquitination 4 h after depolarization [61], which is still consistent with the observations by the Matsuda lab, who showed that Parkin recruitment is not affected by March5 KO after more than 90 min depolarization [60]. Phu et al. later showed that knockdown of March5 reduces overall level of pUb chains after 4 h depolarization, in HeLa cells lacking Parkin [62]. This group also found that the deubiquitinase Usp30 opposes the effect of March5 on the ubiquitination of proteins imported to mitochondria, and Ups30 KO in HEK293 cells increased pUb levels. Thus, March5 at the very least modulates Parkin-dependent mitochondrial protein turnover and may be critical to prime the Parkin-PINK1 pathway in a physiological context.

In conclusion, our work demonstrates that while Parkin does not have intrinsic substrate specificity, it preferably ubiquitinates Mfn2 by virtue of its proximity to PINK1 on the mitochondrial outer membrane. Future work should determine the nature and location of the PINK1:Mfn2 interaction, the mechanism and physiological triggers that lead to the formation of pUb-labeled Mfn2, and the role of March5 in priming the Parkin-PINK1 pathway.

## Acknowledgments

We thank the McGill Pharmacology SPR/MS facility (M. Hancock) and Imaging and Molecular Biology Platform (N. Audet) for assistance with experiments, as well as the Canada Foundation for Innovation for infrastructure support. This work was supported by a Canada Research Chair grant to JFT (#950-229792) and EAF (950-232176), as well as a Canadian Institutes of Health Research (CIHR) grants to KG (FDN-159903), EAF (FDN-154301), and JFT (PJT-153274).

## Materials and Methods

### Cloning and production of purified recombinant proteins

Synthetic codon optimized (DNA Express Inc.) genes of *Rattus norvegicus* Parkin and *Dendroctonus ponderosae* PINK1 (123–570) for *E. coli* expression were cloned into pGEX-6P-1 using BamH1 and Xho1 restriction enzymes. Single-point mutations were introduced using PCR site-directed mutagenesis. Protein expression in BL21 (DE3) *E. coli* cells were induced with 25 µM isopropyl-β-D-thiogalactoside (IPTG) for the different Parkin variants and 100 µM IPTG for DpPINK1. Protein purification was performed as previously described [23]. Briefly, proteins were purified by glutathione-Sepharose agarose affinity and eluted with 20 mM glutathione, followed by 3C cleavage and size-exclusion chromatography on Superdex 200 16/60 (GE Healthcare) in 50 mM Tris-HCl pH 7.5, 120 mM NaCl, and 1 mM dithiothreitol (DTT) or 0.5 mM tris(2-carboxyethyl)phosphine (TCEP). His6-tag UbcH7 was produced in BL21 (DE3) *E. coli* cells transformed with pET28a-LIC-UbcH7 [18]. Protein was purified by Ni-NTA (Qiagen) and eluted with 300 mM Imidazole, followed by gel filtration on Superdex 75 16/60 (GE Healthcare) in 10 mM HEPES pH 7.5, 50 mM NaCl and 1 mM DTT. Protein concentrations were determined with ultraviolet absorption at 280 nm using the theoretical extinction coefficients.

### In vitro phosphorylation of proteins by *Dp*PINK1

Parkin phosphorylation was performed with 3.5 µM GST-*Dp*PINK1 and 20 µM of *Rn*Parkin (in 50 mM Tris-HCl, 120 mM NaCl and 1 mM DTT), 10 mM MgCl_2_ and 1 mM ATP in a reaction volume of 20 µl at 30 °C for 30 to 60 min. GST fused to ubiquitin (GST-Ub) phosphorylation was performed with 3.5 µM *Dp*PINK1 and 43 µM of GST-Ub (in 50 mM Tris-HCl, 120 mM NaCl and 1 mM DTT), 10 mM MgCl_2_ and 1 mM ATP in a reaction volume of 20 µL at 30 °C for 30 to 60 min. Phosphorylated proteins were recovered by gel filtration on Superdex 200 Increase (GE Healthcare) in 50 mM Tris-HCl, 120 mM NaCl pH 7.5. Protein concentrations were determined with ultraviolet absorption at 280 nm using the theoretical extinction coefficients. Phosphorylation was confirmed using intact mass spectrometry (see below).

### Auto-ubiquitination and fluorescent-lysine ubiquitination assays

The auto-ubiquitination assays (Figure 1(c)) were performed at 37°C for 120 min, in 50 mM Tris-HCl pH 7.5, 120 mM NaCl, 1 mM DTT, 2 mM ATP, 50 µM Ub (Boston Biochem), 10 mM MgCl2, 50 nM E1 (Boston Biochem), 2 µM UbcH7 and 1.5 µM GST-*Rn*Parkin. Reactions were stopped with the addition of SDS–PAGE sample buffer containing 50 mM DTT and analysed using SDS-PAGE and CBB staining. For the experiment shown in Figure S2(c), the same conditions were used, but Tris-HCl and NaCl were replaced by mitochondrial isolation buffer (20 mM HEPES/KOH pH 7.4, 220 mM mannitol, 10 mM potassium acetate, 70 mM sucrose), with or without 0.1% NP-40. The ubiquitination assay with fluorescent lysine (Figure 1(d)) was performed at 37°C for 60 min, in 50 mM HEPES pH 7.4, 50 mM NaCl, 0.5 mM DTT, 2 mM ATP, 50 µM Ub, 10 mM MgCl2, 50 nM E1 4 µM UbcH7, 2 µM GST-*Rn*Parkin, and 1 mM 5-FAM-Lysine (Anaspec), prepared from a 20 mM stock in DMSO. Reactions were stopped SDS–PAGE sample buffer containing 50 mM DTT and analysed using SDS-PAGE and fluorescence imaging (Cy2 channel).

### Charging and discharging assays of UbcH7

Ten µg of ubiquitin-conjugating enzyme 2, UbcH7, were charged with 10 µg ubiquitin or N-terminally fluorescein-labelled Ub (FluoUb, Boston Biochem) using 0.02 µg ubiquitin-activating enzyme 1 (His6-E1, Boston Biochem Inc.), 0.5 mM ATP and 10 mM MgCl2 in 50 mM Tris/HCl pH 7.4, 120 mM NaCl and 1 mM TCEP, in a 100 µL reaction. Reactions were performed at 37 °C at different timepoints to a maximum of 60 min.

Ubiquitin-loaded UbcH7 (0.8 µg UbcH7∼Ub or UbcH7∼FluoUb) was mixed with 1.2 µg of *Rn*Parkin (in 50 mM Tris/HCl pH 7.4, 120 mM NaCl and 1 mM TCEP). Mixtures were incubated at 30 °C for up to 40 min to monitor discharging (Figures 2(a,c)). Reactions were stopped with the addition of SDS–PAGE sample buffer containing 12 mM TCEP, resolved on SDS–PAGE and analysed using the Typhoon fluorescent scanner (GE), Coomassie staining, silver staining (Thermo Pierce kit) or transferred to a nitrocellulose membrane and probed with mouse anti-ubiquitin (Covance, 1:10,000) and detected with Clarity Lightning ECL (Bio Rad). Images acquired with an ImageQuant LAS 500 (GE Healthcare). Densitometry and fluorescence intensity analysis was performed using Fiji [63].

### Fluorescent-Ub charging assay on Parkin

Eight µM UbcH7 was charged with 10 µM N-terminal fluorescein-labeled Ub at 37°C for 15 min in 50 mM HEPES pH 7.4, 50 mM NaCl, 0.5 mM TCEP, 4 mM ATP, 20 mM MgCl2, and 100 nM E1 (Figure 2(b)). The master mix was then split and mixed with an equivalent volume of GST-RnParkin WT or mutants (4 µM) and incubated at 37°C for 15 min. Each reaction was split and stopped with SDS–PAGE sample buffer containing either 50 mM TCEP or 100 mM DTT. Reactions were resolved on SDS-PAGE and analyzed using fluorescence imaging (Cy2 channel).

### Ubiquitination of GST-Ub/pUb

Ubiquitination assays (Figure 5(b)) were performed at 37 °C for 30 min, in buffer containing 50 mM Tris-HCl pH 8.0, 5 mM MgCl2, 4 µM UbcH7 or UbcH7∼Ub (charged as described above), 0.1 µM *Rn*Parkin or phosphorylated *Rn*Parkin (Parkin^S65^), 50 µM TCEP and 2 µg of GST-Ub or phosphorylated GST-Ub (GST-pUb^S65^). Reactions were stopped with the addition of SDS–PAGE sample buffer containing 100 mM DTT and analysed using gel electrophoresis and CBB staining or immunoblotting. The same reactions (conditions in lanes 9-12 in Figure 5(b)) were repeated and frozen at −80 °C for digestion and mass spectrometry (Figure 5(c) or SDS-PAGE analysis (Figure S3(c)).

### Cell culture, mammalian plasmid DNA, and generation of stable cell lines

U2OS and HeLa cell lines were maintained at 37 °C and 5% CO2 in Dulbecco’s modified Eagle’s medium (DMEM) with 10% fetal bovine serum, 2 mM glutamine, 0.1% penicillin, and 0.1% streptomycin (Wisent Life Sciences). Parkin mutant plasmids were cloned beginning with a Homo sapiens wild type eGFP-parkin vector [36] and performing PCR point mutagenesis (Quikchange Mutagenesis kit: Stratagene) utilizing primers ordered from Invitrogen. Mutations were verified by sequencing at the Genome Québec Facility.

Plasmid DNA was introduced into U2OS cells following a protocol for jetPrime® transfection reagent (Polyplus). Nucleic acid was first diluted in jetPrime® buffer (Polyplus) and combined with jetPrime® transfection reagent before being added to U2OS cells immersed in fresh DMEM. Cells were incubated overnight in the transfection mix and allowed to express eGFP-parkin constructs, a process that was verified by fluorescent light microscopy. Selection of cells stably expressing eGFP-parkin wild type and mutant constructs was accomplished by maintaining them in Dulbecco’s Modified Eagle Medium (DMEM) laced with 200 mg/L G418 (Multicell). U2OS cells were grown in G418-DMEM for a period of ∼2 weeks and passaged using trypsin as necessary.

### Mitochondrial GFP-parkin recruitment and microscopy

DMEM containing CCCP at a final concentration of 20 μM, or 01% DMSO, was added to plates growing GFP-Parkin-expressing U2OS cells for 1, 2, 4 or 24 h. For immunoblots, cells were harvested on ice in cold phosphate buffered saline (PBS). Cells were washed and lysed in RIPA buffer with a protease inhibitor cocktail (1:100 benzamidine, 1:10000 aprotinin, 1:10000 leupeptin, 1:500 PMSF). Protein concentration in these samples was quantified using the Pierce BCA Assay (Thermo Scientific). Samples were mixed with SDS-PAGE sample buffer with 100 mM DTT and resolved by SDS-PAGE. Proteins were transferred to a nitrocellulose membrane and stained with ponceau. After staining, membranes were washed with PBS-0.1% Tween 20 (PBS-T), blocked with 5% milk/PBS-T, and incubated with primary antibodies overnight. Primaries used in all experiments included mouse anti-Parkin (1:20000; Santa Cruz), rabbit anti-Mfn1 and anti-Mfn2 (1:2000, Santa Cruz), and mouse anti-VDAC1 (1:10000, Abcam) diluted in PBS-T with 3% BSA. Bound protein was detected on film using HRP-coupled secondary antibodies (donkey- or goat-anti-rabbit/anti-mouse), treated with Western Lightning ECL/Super ECL as needed (Perkin Elmer).

For microscopy. cells grown on coverslips were fixed with 4% formaldehyde-PBS (prepared from 16% formaldehyde stock, Thermo Fisher Scientific) for 15 min at 37 °C. After fixation, cells were washed four times with 1× PBS, permeabilized with 1× PBS containing 0.25% Triton X-100 for 10 min, and blocked with 3% BSA + PBST (1× PBS + 0.1% Tween-20) for 1 h at RT. Cells were incubated overnight at 4 °C with rabbit anti-Tom20 (1:1000, Santa Cruz) followed by Alexa Fluor® 555 donkey anti-rabbit (Invitrogen). Cells were washed three times in PBS, and coverslips were mounted with fluorescent mounting medium (Dako). Confocal images were acquired on an LSM 510 Meta confocal microscope (Zeiss) using a 25×, 0.8NA or 63×, 1.4 NA objective. Excitation wavelengths of 488 nm (GFP-Parkin) and 633 nm (Tom20) were used. For quantification, slides were blinded for three independent experiments and 100 cells were counted per experiment. Cells were analyzed for the number of cells with TOM20 staining (CCCP 24 h), the number of cells with GFP-parkin co-localization on TOM20-positive mitochondria (CCCP 1–2 h), or for the number of cells containing GFP-parkin puncta or aggregates co-localizing with TOM20-positive mitochondria (CCCP 24 h).

### Mitochondria isolation and *in organello* ubiquitination assay

HeLa cells treated with 10 μM CCCP or DMSO for 3 h were suspended in mitochondrial isolation buffer (20 mM HEPES/KOH pH 7.4, 220 mM mannitol, 10 mM potassium acetate, 70 mM sucrose) on ice. Cells were disrupted by nitrogen cavitation and cell homogenates were centrifuged at 600g for 5 min at 4 °C to obtain a post-nuclear supernatant. Cytosolic fractions were collected by two further centrifugation steps for 10 min at 4 °C, the first at 10,000g and the second at 12,000g.

Mitochondria pellets were suspended in mitochondria isolation buffer to a concentration of 2 mg/mL and stored at −80 °C until further use. Forty micrograms of CCCP- or DMSO-treated mitochondria were supplemented with a ubiquitination reaction mix (20 nM E1, 100 nM of UbcH7, 5 μM ubiquitin, 1 mM ATP, 5 mM MgCl2 and 50 μM TCEP in mitochondria isolation buffer) and 100 nM of recombinant *Rn*Parkin in a 40 μL reaction. After a 30-min incubation at 37°C reactions were stopped with 3X sample buffer with 100 mM DTT and analyzed by western-blotting. Reactions were loaded on 8% tris-glycine. Proteins were transferred to nitrocellulose and stained with Ponceau. Membranes were blocked with 5% milk in PBS-T (0.1% Tween 20) and incubated with rabbit anti-mitofusin 2 (mAb D2D10, Cell Signaling), rabbit anti-mitofusin 1 (mAb 13196S, Cell Signaling), rabbit anti-Miro1 (mAb 14016S, Cell Signaling), rabbit anti-HK1 (mAb C35C4, Cell Signaling), rabbit anti-HK2 (mAb C64G5, Cell Signaling), rabbit anti-TOM20 (pAb FL-145, Santa Cruz Biotechnology), rabbit anti-TOM70 (pAb C-18, Santa Cruz Biotechnology), mouse anti-Parkin (mAb Prk8, Cell Signaling), rabbit anti-PINK1 (mAb D8G3, Cell Signaling), mouse anti-Ub (mAb P4D1, Cell Signaling), rabbit anti-phospho ubiquitin S65 (Millipore), rabbit anti-PDH (mAb C54G1, Cell Signaling), or rabbit anti-VDAC (mAb D73D12, Cell Signaling) diluted in PBS-T with 3% bovine serum albumin (BSA). Membranes were washed with PBS-T and incubated with HRP-coupled goat anti-mouse or anti-rabbit IgG antibodies (Cell Signaling). Detection was performed with Clarity Lightning ECL (Bio Rad) and images acquired with an ImageQuant LAS 500 (GE Healthcare).

### Proximity Ligation Assay for Proteins Interaction Studies

Protein-protein interactions were analyzed using Duolink in situ orange starter fluorescence kit (mouse/rabbit, Sigma-Aldrich) in human osteosarcoma U2OS cells (WT, Mfn2 KO and PINK1 KO). Cells were grown to confluency on coverslips (Fisherbrand) and 5 μg pCMV(d1) TNT PINK1(WT)-3HA plasmids, obtained from Noriyuki Matsuda for attenuated PINK1 expression, were transfected using Lipofectamine 3000 Reagent (ThermoFisher Scientific). Cells were grown for an additional 48 hours post transfection. Cells were treated with 10 μM CCCP or DMSO for 3 hours before fixation and permeabilization using 4% PFA/0.1% Triton-X-100 for 10 min at room temperature. Samples were blocked using Duolink blocking solution in a preheated humidified chamber at 37 °C for one hour. Primary antibody solution mix contains two primary antibodies raised in two different species (mouse and rabbit) targeting the proteins of interest: rabbit anti-mitofusin 2 (1:50, mAb D2D10, Cell Signaling), rabbit anti-VDAC (1:200, mAb D73D12, Cell Signaling), mouse anti-HA (1:100, mAb D73D12, Cell Signaling), rabbit anti-mitofusin 1 (1:100, mAb 13196S, Cell Signaling), rabbit anti-Miro1 (1:50, mAb 14016S, Cell Signaling), rabbit anti-pUb (1:100, mAb 37642S, Cell Signaling), or mouse anti-mitofusin 2 (1:50, mAb 661-757, Abnova) was added for overnight incubation at 4 °C. The next day, cells were incubated with proximity ligation assay probes PLUS and MINUS diluted 1:5 for one hour at 37 °C. After ligation, the samples were incubated with amplification polymerase solution for 100 min at 37 °C, protected from light. The endoplasmic reticulum was stained using an Alexa488-coupled anti-calnexin (1:200, mAb AF18, ThermoFisher Scientific) in 10% goat serum overnight at 4 °C. Cell nuclei were stained using DAPI (1:1000, ThermoFisher Scientific) for 10 min at room temperature. All samples were mounted with mounting medium (Dako, Agilent). Images were acquired with a 40x Plan Apo oil-immersion objective using a TCS SP8 confocal microscope (Leica).

### Protein digestion and mass spectrometry

Ubiquitination reaction samples (3 µg protein) were diluted in denaturing buffer (3 M urea, 25 mM TEAB pH 8.5, 0.5 mM EDTA) and reduced using 2 mM TCEP for 10 min at 37 °C, followed by alkylation with 50 mM chloroacetamide for 30 min at room temperature in the dark. Chloroacetamide was used to avoid iodoacetamide-induced artefacts that mimic ubiquitination [64]. Samples were diluted with 50 mM TEAB pH 8.5 to 1 M urea and digested with 0.5 µg trypsin (Sigma) for 3 h at 37 °C. Digested peptides were purified using C18 Spin Columns (ThermoFisher) and resuspended in 0.1% formic acid. Peptides (0.5 µg) were captured and eluted from an Acclaim PepMap100 C18 column with a 2 h gradient of acetonitrile in 0.1% formic acid at 200 nl/min. The eluted peptides were analysed with an Impact II Q-TOF spectrometer equipped with a Captive Spray nanoelectrospray source (Bruker). Data were acquired using data-dependent automatic tandem mass spectrometry (auto-MS/MS) and analysed with MaxQuant using a standard search procedure against a custom-made FASTA file including GST-Ub, Parkin, Ub and UbcH7. Methionine oxidation, Lys ubiquitination (diGly) and Ser/Thr phosphorylation were included as variable modifications. Cysteine carbamylation was included as fixed modification.

## Author Contributions

J.-F.T. conceived experiments, performed biochemical assays and wrote the manuscript. M.V. performed protein purification, biochemical experiments and *in organello* assays, and wrote the manuscript. Y.L. performed protein purification, *in organello* and PLA assays. V.S. and N.C. performed protein purification. S.R. performed in organello assays and phosphorylation assays. T.M.D. & J.D.K. performed cell-based mitophagy assays. K.G. and E. A. Fon conceived experiments and edited the manuscript.

## Figures and Figure legends

**Figure S1.**
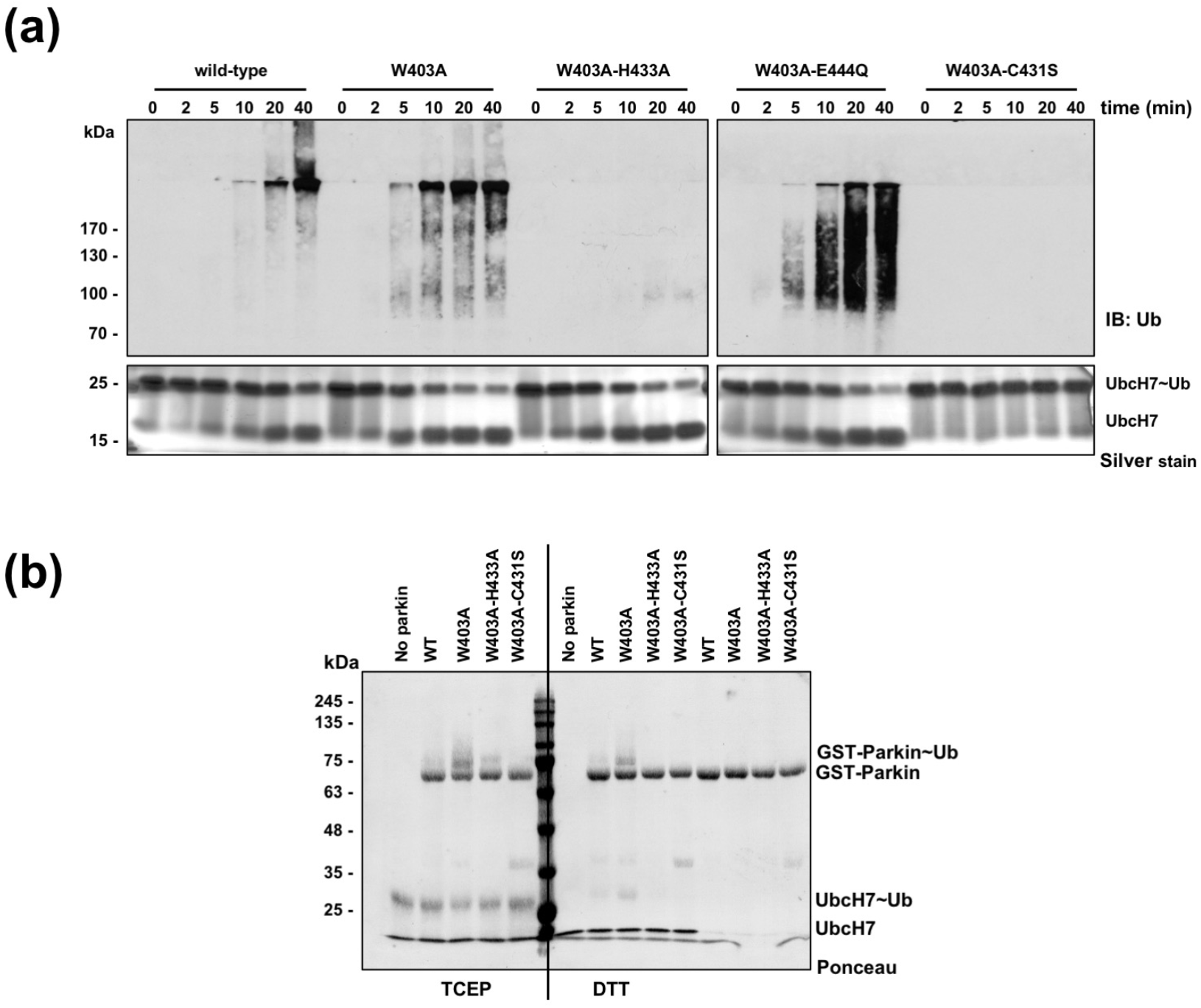
His433 is required for the E3 ubiquitin ligase activity of parkin. **(a)** Immunoblot (top) and silver-stained (bottom) SDS-PAGE of UbcH7∼Ub discharge assays with GST-parkin mutants. Reactions were stopped with sample buffer containing tris(2-carboxyethyl)phosphine (TCEP) to reduce disulphide bonds but keep thioester bonds intact. Data for WT, W403A and C431S were originally reported in Trempe et al. (2013) **(b)** Ponceau-stained transferred membrane showing Ub-linkage after mixing non-labelled Ub, E1, ATP and UbcH7 with GST-parkin. Incubation mixes stopped with TCEP or DTT sample buffer. The last four lanes are reaction controls without UbcH7.

**Figure S2.**
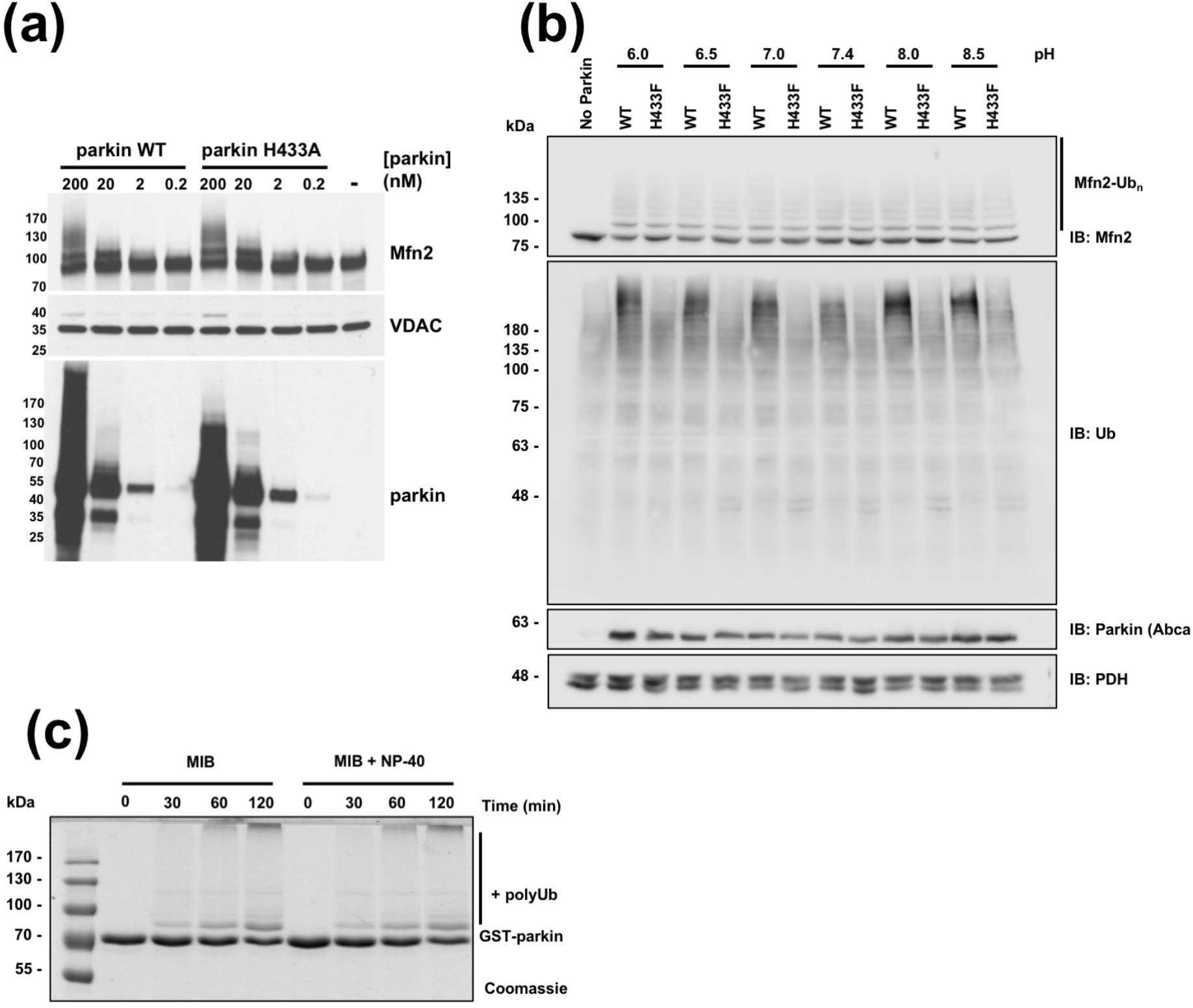
Mutation of Parkin His433 has differential effects on substrate ubiquitination. **(a)** Immunoblot of *in organello* ubiquitination assay showing that ubiquitination of Mfn2 is not affected by H433A mutation and is proportional to the amount of Parkin in the reaction. Positive ubiquitination is detected by high-molecular weight bands above the unmodified protein bands. **(b)** Immunoblot of *in organello* ubiquitination assay showing that the overall ubiquitination levels detected at high molecular weight protein bands is significantly affected when His433 is mutated, whereas Mfn2 ubiquitination is not. **(c)** CBB-stained SDS-PAGE of Parkin auto*-*ubiquitination assay in the presence of the non-ionic detergent NP40.

**Figure S3.**
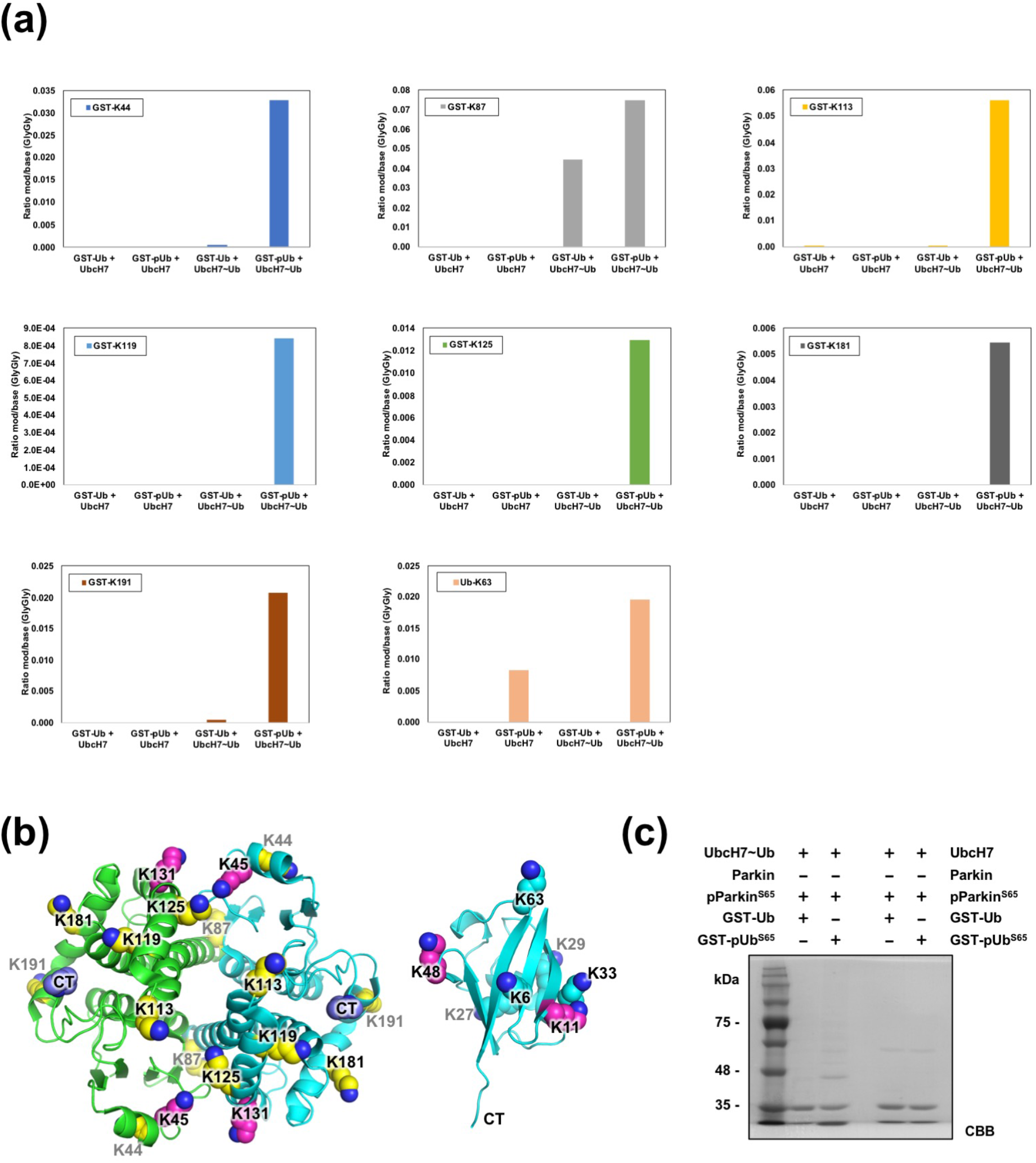
Parkin preferably ubiquitinates substrates tethered to phospho-ubiquitin. **(a)** Mass spectrometry analysis of the ubiquitination sites found after ubiquitination assay reactions where Parkin (phospho or non phospho) was incubated with UbcH7 or UbcH7∼Ub and GST-Ub or GST-pUb^S65^. **(b)** 3D structure of glutathione S-transferase (GST; PDB 6RWD, left) and ubiquitin (PDB 1UBQ, right). Target lysine side-chains are shown as spheres; major sites in magenta, minor sites in yellow. **(c)** CBB-stained SDS-PAGE of the ubiquitination assay where purified Parkin or phosphorylated Parkin (pParkin^S65^) was incubated with UbcH7∼Ub as a ubiquitin source and GST-Ub or GST-pUb^S65^ as the ubiquitin acceptor. Right lanes with UbcH7 only are reaction controls.

**Figure S4:**
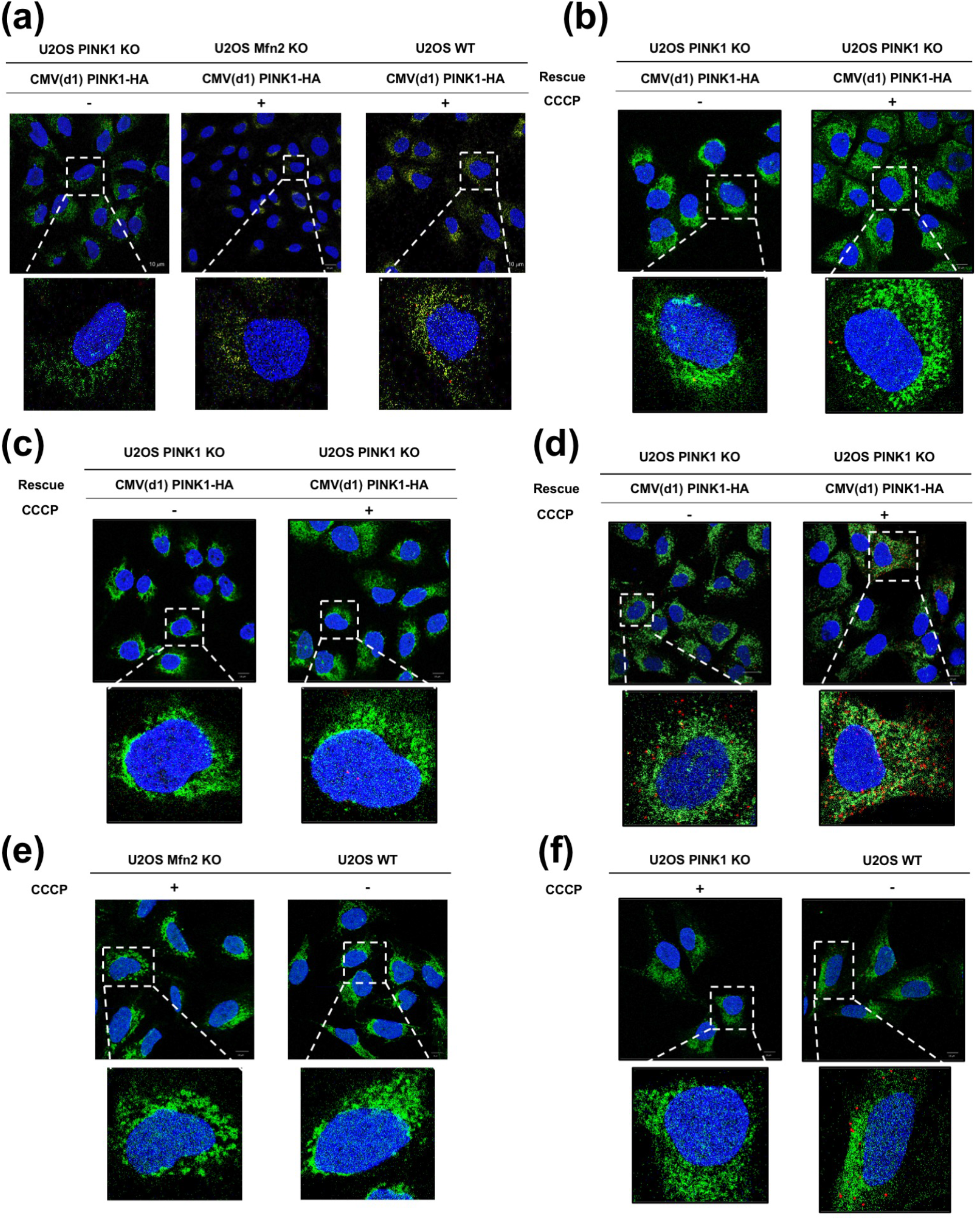
Proximity ligation assay confocal microscopy images to assess the co-localization of PINK1 and outer mitochondria membrane proteins on damaged mitochondria. **(a)** Experiment controls for PLA targeting Mfn2 and PINK1 including U2OS WT, PINK1 and Mfn2 KO cell lines expressing recombinant PINK1-HA. PLA targeting Mfn1 and PINK1 **(b)**, Miro and PINK1 **(c)**, VDAC and PINK1 **(d)** and targeting Mfn2 and pUb **(e)** in U2OS knockout cell line backgrounds.

## References and Notes

[1] Kitada T, Asakawa S, Hattori N, Matsumine H, Yamamura Y, Minoshima S, et al. Mutations in the parkin gene cause autosomal recessive juvenile parkinsonism. Nature. 1998;392:605–8.

[2] Winklhofer KF. Parkin and mitochondrial quality control: toward assembling the puzzle. Trends Cell Biol. 2014;24:332–41.

[3] Valente EM, Abou-Sleiman PM, Caputo V, Muqit MM, Harvey K, Gispert S, et al. Hereditary early-onset Parkinson’s disease caused by mutations in PINK1. Science. 2004;304:1158–60.

[4] Koyano F, Okatsu K, Kosako H, Tamura Y, Go E, Kimura M, et al. Ubiquitin is phosphorylated by PINK1 to activate parkin. Nature. 2014;510:162–6.

[5] Kane LA, Lazarou M, Fogel AI, Li Y, Yamano K, Sarraf SA, et al. PINK1 phosphorylates ubiquitin to activate Parkin E3 ubiquitin ligase activity. The Journal of cell biology. 2014;205:143–53.

[6] Kondapalli C, Kazlauskaite A, Zhang N, Woodroof HI, Campbell DG, Gourlay R, et al. PINK1 is activated by mitochondrial membrane potential depolarization and stimulates Parkin E3 ligase activity by phosphorylating Serine 65. Open Biol. 2012;2:120080.

[7] Narendra DP, Jin SM, Tanaka A, Suen DF, Gautier CA, Shen J, et al. PINK1 is selectively stabilized on impaired mitochondria to activate Parkin. PLoS Biol. 2010;8:e1000298.

[8] Matsuda N, Sato S, Shiba K, Okatsu K, Saisho K, Gautier CA, et al. PINK1 stabilized by mitochondrial depolarization recruits Parkin to damaged mitochondria and activates latent Parkin for mitophagy. J Cell Biol. 2010;189:211–21.

[9] Geisler S, Holmstrom KM, Skujat D, Fiesel FC, Rothfuss OC, Kahle PJ, et al. PINK1/Parkin-mediated mitophagy is dependent on VDAC1 and p62/SQSTM1. Nature cell biology. 2010;12:119–31.

[10] Narendra D, Tanaka A, Suen DF, Youle RJ. Parkin is recruited selectively to impaired mitochondria and promotes their autophagy. J Cell Biol. 2008;183:795–803.

[11] Sarraf SA, Raman M, Guarani-Pereira V, Sowa ME, Huttlin EL, Gygi SP, et al. Landscape of the PARKIN-dependent ubiquitylome in response to mitochondrial depolarization. Nature. 2013;496:372–6.

[12] Chan NC, Salazar AM, Pham AH, Sweredoski MJ, Kolawa NJ, Graham RL, et al. Broad activation of the ubiquitin-proteasome system by Parkin is critical for mitophagy. Hum Mol Genet. 2011;20:1726–37.

[13] Tanaka A, Cleland MM, Xu S, Narendra DP, Suen DF, Karbowski M, et al. Proteasome and p97 mediate mitophagy and degradation of mitofusins induced by Parkin. The Journal of cell biology. 2010;191:1367–80.

[14] Ordureau A, Heo JM, Duda DM, Paulo JA, Olszewski JL, Yanishevski D, et al. Defining roles of PARKIN and ubiquitin phosphorylation by PINK1 in mitochondrial quality control using a ubiquitin replacement strategy. Proceedings of the National Academy of Sciences of the United States of America. 2015;112:6637–42.

[15] Lazarou M, Sliter DA, Kane LA, Sarraf SA, Wang C, Burman JL, et al. The ubiquitin kinase PINK1 recruits autophagy receptors to induce mitophagy. Nature. 2015;524:309–14.

[16] McLelland GL, Soubannier V, Chen CX, McBride HM, Fon EA. Parkin and PINK1 function in a vesicular trafficking pathway regulating mitochondrial quality control. The EMBO journal. 2014;33:282–95.

[17] Wenzel DM, Lissounov A, Brzovic PS, Klevit RE. UBCH7 reactivity profile reveals parkin and HHARI to be RING/HECT hybrids. Nature. 2011;474:105–8.

[18] Trempe JF, Sauvé V, Grenier K, Seirafi M, Tang MY, Menade M, et al. Structure of parkin reveals mechanisms for ubiquitin ligase activation. Science. 2013;340:1451–5.

[19] Wauer T, Komander D. Structure of the human Parkin ligase domain in an autoinhibited state. The EMBO journal. 2013;32:2099–112.

[20] Riley BE, Lougheed JC, Callaway K, Velasquez M, Brecht E, Nguyen L, et al. Structure and function of Parkin E3 ubiquitin ligase reveals aspects of RING and HECT ligases. Nat Commun. 2013;4:1982.

[21] Chaugule VK, Burchell L, Barber KR, Sidhu A, Leslie SJ, Shaw GS, et al. Autoregulation of Parkin activity through its ubiquitin-like domain. EMBO J. 2011;30:2853–67.

[22] Wauer T, Simicek M, Schubert A, Komander D. Mechanism of phospho-ubiquitin-induced PARKIN activation. Nature. 2015;524:370–4.

[23] Sauvé V, Lilov A, Seirafi M, Vranas M, Rasool S, Kozlov G, et al. A Ubl/ubiquitin switch in the activation of Parkin. The EMBO journal. 2015;34:2492–505.

[24] Kumar A, Aguirre JD, Condos TE, Martinez-Torres RJ, Chaugule VK, Toth R, et al. Disruption of the autoinhibited state primes the E3 ligase parkin for activation and catalysis. The EMBO journal. 2015;34:2506–21.

[25] Kazlauskaite A, Martinez-Torres RJ, Wilkie S, Kumar A, Peltier J, Gonzalez A, et al. Binding to serine 65-phosphorylated ubiquitin primes Parkin for optimal PINK1-dependent phosphorylation and activation. EMBO Rep. 2015;16:939–54.

[26] Tang MY, Vranas M, Krahn AI, Pundlik S, Trempe JF, Fon EA. Structure-guided mutagenesis reveals a hierarchical mechanism of Parkin activation. Nat Commun. 2017;8:14697.

[27] Rasool S, Soya N, Truong L, Croteau N, Lukacs GL, Trempe JF. PINK1 autophosphorylation is required for ubiquitin recognition. EMBO Rep. 2018;19:e44981.

[28] Sauvé V, Sung G, Soya N, Kozlov G, Blaimschein N, Miotto L, et al. Mechanism of parkin activation by phosphorylation. Nat Struct Mol Biol. 2018;25:623–30.

[29] Gladkova C, Maslen SL, Skehel JM, Komander D. Mechanism of parkin activation by PINK1. Nature. 2018;559:410–4.

[30] Matsuda N, Kitami T, Suzuki T, Mizuno Y, Hattori N, Tanaka K. Diverse effects of pathogenic mutations of Parkin that catalyze multiple monoubiquitylation in vitro. J Biol Chem. 2006;281:3204–9.

[31] Hampe C, Ardila-Osorio H, Fournier M, Brice A, Corti O. Biochemical analysis of Parkinson’s disease-causing variants of Parkin, an E3 ubiquitin-protein ligase with monoubiquitylation capacity. Hum Mol Genet. 2006;15:2059–75.

[32] Pahari S, Sun L, Alexov E. PKAD: a database of experimentally measured pKa values of ionizable groups in proteins. Database (Oxford). 2019;2019.

[33] Spratt DE, Julio Martinez-Torres R, Noh YJ, Mercier P, Manczyk N, Barber KR, et al. A molecular explanation for the recessive nature of parkin-linked Parkinson’s disease. Nat Commun. 2013;4:1983.

[34] Stieglitz B, Rana RR, Koliopoulos MG, Morris-Davies AC, Schaeffer V, Christodoulou E, et al. Structural basis for ligase-specific conjugation of linear ubiquitin chains by HOIP. Nature. 2013;503:422–6.

[35] Ordureau A, Sarraf SA, Duda DM, Heo JM, Jedrykowski MP, Sviderskiy VO, et al. Quantitative Proteomics Reveal a Feedforward Mechanism for Mitochondrial PARKIN Translocation and Ubiquitin Chain Synthesis. Mol Cell. 2014;56:360–75.

[36] Durcan TM, Tang MY, Perusse JR, Dashti EA, Aguileta MA, McLelland GL, et al. USP8 regulates mitophagy by removing K6-linked ubiquitin conjugates from parkin. The EMBO journal. 2014;33:2473–91.

[37] Ordureau A, Paulo JA, Zhang W, Ahfeldt T, Zhang J, Cohn EF, et al. Dynamics of PARKIN-Dependent Mitochondrial Ubiquitylation in Induced Neurons and Model Systems Revealed by Digital Snapshot Proteomics. Mol Cell. 2018;70:211–27.

[38] McLelland GL, Goiran T, Yi W, Dorval G, Chen CX, Lauinger ND, et al. Mfn2 ubiquitination by PINK1/parkin gates the p97-dependent release of ER from mitochondria to drive mitophagy. eLife. 2018;7:e32866.

[39] Rakovic A, Shurkewitsch K, Seibler P, Grunewald A, Zanon A, Hagenah J, et al. Phosphatase and tensin homolog (PTEN)-induced putative kinase 1 (PINK1)-dependent ubiquitination of endogenous Parkin attenuates mitophagy: study in human primary fibroblasts and induced pluripotent stem cell-derived neurons. The Journal of biological chemistry. 2013;288:2223–37.

[40] Yi W, MacDougall EJ, Tang MY, Krahn AI, Gan-Or Z, Trempe JF, et al. The landscape of Parkin variants reveals pathogenic mechanisms and therapeutic targets in Parkinson’s disease. Human molecular genetics. 2019;28:2811–25.

[41] Roberts RF, Wade-Martins R, Alegre-Abarrategui J. Direct visualization of alpha-synuclein oligomers reveals previously undetected pathology in Parkinson’s disease brain. Brain. 2015;138:1642–57.

[42] Okatsu K, Oka T, Iguchi M, Imamura K, Kosako H, Tani N, et al. PINK1 autophosphorylation upon membrane potential dissipation is essential for Parkin recruitment to damaged mitochondria. Nat Commun. 2012;3:1016.

[43] Smit JJ, Monteferrario D, Noordermeer SM, van Dijk WJ, van der Reijden BA, Sixma TK. The E3 ligase HOIP specifies linear ubiquitin chain assembly through its RING-IBR-RING domain and the unique LDD extension. EMBO J. 2012;31:3833–44.

[44] Lechtenberg BC, Rajput A, Sanishvili R, Dobaczewska MK, Ware CF, Mace PD, et al. Structure of a HOIP/E2∼ubiquitin complex reveals RBR E3 ligase mechanism and regulation. Nature. 2016;529:546–50.

[45] Stieglitz B, Morris-Davies AC, Koliopoulos MG, Christodoulou E, Rittinger K. LUBAC synthesizes linear ubiquitin chains via a thioester intermediate. EMBO Rep. 2012;13:840–6.

[46] Kelsall IR, Zhang J, Knebel A, Arthur JSC, Cohen P. The E3 ligase HOIL-1 catalyses ester bond formation between ubiquitin and components of the Myddosome in mammalian cells. Proceedings of the National Academy of Sciences of the United States of America. 2019;116:13293–8.

[47] Seenivasan R, Hermanns T, Blyszcz T, Lammers M, Praefcke GJK, Hofmann K. Mechanism and chain specificity of RNF216/TRIAD3, the ubiquitin ligase mutated in Gordon Holmes syndrome. Human molecular genetics. 2019;28:2862–73.

[48] Michel MA, Swatek KN, Hospenthal MK, Komander D. Ubiquitin Linkage-Specific Affimers Reveal Insights into K6-Linked Ubiquitin Signaling. Mol Cell. 2017;68:233–46 e5.

[49] de Brito OM, Scorrano L. Mitofusin 2 tethers endoplasmic reticulum to mitochondria. Nature. 2008;456:605–10.

[50] Basso V, Marchesan E, Peggion C, Chakraborty J, von Stockum S, Giacomello M, et al. Regulation of Endoplasmic Reticulum-Mitochondria contacts by Parkin via Mfn2. Pharmacological research : the official journal of the Italian Pharmacological Society. 2018;138:43–56.

[51] Gelmetti V, De Rosa P, Torosantucci L, Marini ES, Romagnoli A, Di Rienzo M, et al. PINK1 and BECN1 relocalize at mitochondria-associated membranes during mitophagy and promote ER-mitochondria tethering and autophagosome formation. Autophagy. 2017;13:654–69.

[52] Gandhi S, Wood-Kaczmar A, Yao Z, Plun-Favreau H, Deas E, Klupsch K, et al. PINK1-associated Parkinson’s disease is caused by neuronal vulnerability to calcium-induced cell death. Molecular cell. 2009;33:627–38.

[53] Friedman JR, Lackner LL, West M, DiBenedetto JR, Nunnari J, Voeltz GK. ER tubules mark sites of mitochondrial division. Science. 2011;334:358–62.

[54] Kleele T, Rey T, Winter J, Zaganelli S, Mahecic D, Perreten Lambert H, et al. Distinct fission signatures predict mitochondrial degradation or biogenesis. Nature. 2021;593:435–9.

[55] Di Rita A, Peschiaroli A, P Da, Strobbe D, Hu Z, Gruber J, et al. HUWE1 E3 ligase promotes PINK1/PARKIN-independent mitophagy by regulating AMBRA1 activation via IKKalpha. Nat Commun. 2018;9:3755.

[56] Yun J, Puri R, Yang H, Lizzio MA, Wu C, Sheng ZH, et al. MUL1 acts in parallel to the PINK1/parkin pathway in regulating mitofusin and compensates for loss of PINK1/parkin. eLife. 2014;3:e01958.

[57] Braschi E, Zunino R, McBride HM. MAPL is a new mitochondrial SUMO E3 ligase that regulates mitochondrial fission. EMBO Rep. 2009;10:748–54.

[58] Yonashiro R, Ishido S, Kyo S, Fukuda T, Goto E, Matsuki Y, et al. A novel mitochondrial ubiquitin ligase plays a critical role in mitochondrial dynamics. The EMBO journal. 2006;25:3618–26.

[59] Nakamura N, Kimura Y, Tokuda M, Honda S, Hirose S. MARCH-V is a novel mitofusin 2-and Drp1-binding protein able to change mitochondrial morphology. EMBO Rep. 2006;7:1019–22.

[60] Koyano F, Yamano K, Kosako H, Tanaka K, Matsuda N. Parkin recruitment to impaired mitochondria for nonselective ubiquitylation is facilitated by MITOL. The Journal of biological chemistry. 2019;294:10300–14.

[61] Ordureau A, Paulo JA, Zhang J, An H, Swatek KN, Cannon JR, et al. Global Landscape and Dynamics of Parkin and USP30-Dependent Ubiquitylomes in iNeurons during Mitophagic Signaling. Mol Cell. 2020;77:1124–42 e10.

[62] Phu L, Rose CM, Tea JS, Wall CE, Verschueren E, Cheung TK, et al. Dynamic Regulation of Mitochondrial Import by the Ubiquitin System. Mol Cell. 2020;77:1107–23 e10.

[63] Schindelin J, Arganda-Carreras I, Frise E, Kaynig V, Longair M, Pietzsch T, et al. Fiji: an open-source platform for biological-image analysis. Nat Methods. 2012;9:676–82.

[64] Nielsen ML, Vermeulen M, Bonaldi T, Cox J, Moroder L, Mann M. Iodoacetamide-induced artifact mimics ubiquitination in mass spectrometry. Nat Methods. 2008;5:459–60.

